# Reconstitution of human kinetochore in mitotic cell extracts reveals permitted and restricted assembly steps

**DOI:** 10.1101/2020.07.15.203109

**Authors:** Ekaterina V. Tarasovetc, Praveen Kumar Allu, Iain M. Cheeseman, Ben E. Black, Ekaterina L. Grishchuk

**Affiliations:** Department of Physiology, Perelman School of Medicine, University of Pennsylvania, Philadelphia, PA, USA, 19104; Department of Biochemistry and Biophysics, Perelman School of Medicine, University of Pennsylvania, Philadelphia PA, USA, 19104; Whitehead Institute for Biomedical Research, 455 Main Street, Cambridge, MA, USA, 02142

## Abstract

Assembly of a functional kinetochore is critical for accurate chromosome segregation. Hierarchical recruitment of soluble components during kinetochore assembly is a highly regulated mitotic event, but the underlying steps are not well understood. In yeast and *Xenopus* egg extracts, soluble kinetochore components can spontaneously assemble into microtubule-binding subcomplexes. Although the molecular interactions among specific kinetochore components are evolutionary conserved in eukaryotes, it remains unclear which *de novo* assembly steps are permitted in extracts of mitotic human cells. By analyzing the recruitment of GFP-fused kinetochore proteins from human mitotic cell extracts to inner kinetochore components immobilized on microbeads, we reconstructed the interaction between CENP-C and CENP-A–containing nucleosomes. However, subsequent phospho-dependent binding of the Mis12 complex was less efficient, whereas binding of the Ndc80 complex was inhibited. Consistently, the microtubule-binding activity of native kinetochore components, as well as those assembled using a combination of native and recombinant human proteins, was weaker than that of recombinant Ndc80 complex alone. Such inhibitory mechanisms that prevent interactions between different kinetochore components are likely to guard against spurious formation of kinetochores in the cytosol of mitotic human cells, and imply existence of specific regulatory mechanisms that permit these interactions at the assembling kinetochore.

## Introduction

Accurate chromosome segregation depends on proper interactions between spindle microtubules and the kinetochore, a multiprotein complex located on each centromere. Kinetochore assembly is a complex process, requiring the execution of multiple binding reactions in a specific sequence at a dedicated chromosomal location (Cheeseman 2014, Nagpal et al. 2016, Musacchio et al. 2017). Kinetochore assembly is nucleated by inner kinetochore proteins localized at the centromere, which is marked by centromere protein A (CENP-A) containing nucleosomes (Fukagawa et al. 2014, McKinley et al. 2016). At the onset of mitosis, the CENP-A nucleosomes and constitutive centromere-associated protein network (CCAN) recruit multiple copies of outer kinetochore proteins from their soluble pools. Among them are the Ndc80 complex, Mis12 complex, and Knl1 protein, constituting the KMN network which links centromeres and spindle microtubules (Cheeseman 2014, Nagpal et al. 2016, Musacchio et al. 2017). Ndc80 complex, which consists of four subunits (Ndc80/Hec1, Nuf2, Spc24, and Spc25) is the major microtubule-binding component of the kinetochore (reviewed in (Cheeseman et al. 2006, Musacchio et al. 2017)). Previous work identified critical molecular interfaces that mediate interactions between Ndc80 complex and other KMN component Mis12 complex (Figure 1A). The Ndc80 complex, which has reduced microtubule-binding affinity on its own, binds to the Mis12 complex via the Spc24/Spc25 subunits, and this interaction alleviates the intra-molecular inhibition (Cheeseman et al. 2006, Kudalkar et al. 2015, Scarborough et al. 2019). The Mis12 complex, consisting of four subunits (Mis12, Pmf1, Nsl1 and Dsn1), binds to the Ndc80 complex via the C-termini of Dsn1 and Nsl1 subunits (Petrovic et al. 2010, Petrovic et al. 2016).

**Figure 1.**
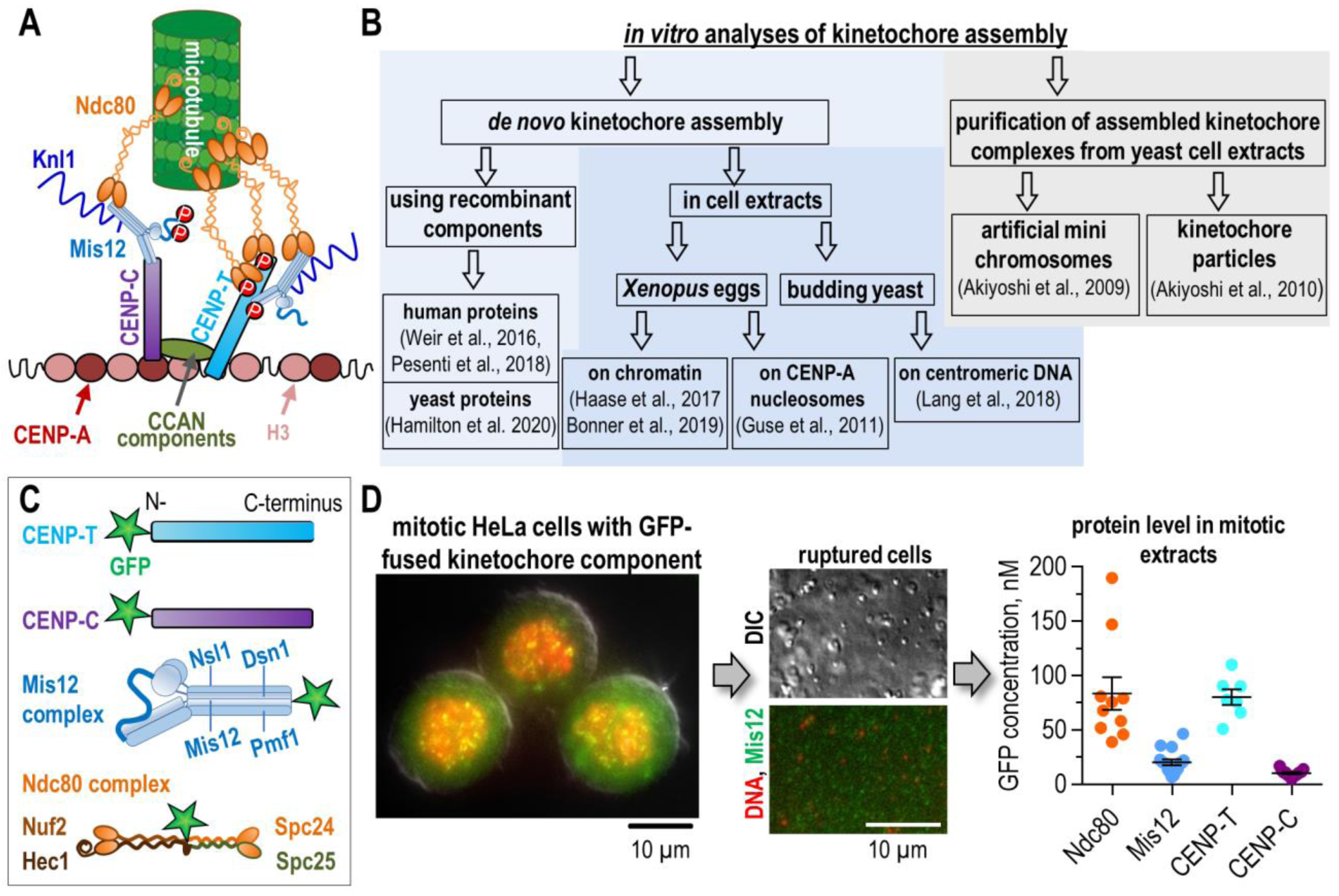
Strategies for reconstructing kinetochores and our experimental approach. (A) Principal architecture of mitotic kinetochore and its binding to microtubules; see text for details. CCAN, constitutive centromere associated network. Letters “P” on CENP-T indicate phosphorylation-dependent activation of its binding to Ndc80 and Mis12 complexes; letters “P” on Mis12 indicate phosphorylation-dependent activation of its binding to CENP-C. (B) Previous experimental approaches to reconstitute kinetochore assembly and function *in vitro*; see text for additional references. (C) Schematic of GFP-fused proteins stably expressed in HeLa cells. Experiments with Ndc80 complexes used three different cell lines: Nuf2-GFP (shown), GFP-Spc24, and GFP-Spc25. (D) Key steps of our experimental approach. Left image shows representative HeLa cell that was fixed and stained with propidium iodide to reveal DNA; green signal is from Mis12-GFP. Ruptured cells (images in the middle) were centrifuged to remove insoluble cellular components. Graph shows concentration of GFP-fused kinetochore proteins in clarified cell extracts. Each colored point represents average bead brightness from independent experiments, during which 50–100 beads were analyzed. For more detailed statistics, see Source data. Black lines show mean with standard error of the mean (SEM). The column for native Ndc80-GFP combines data from cell lines expressing GFP-fused Nuf2, Spc24, and Spc25.

In human cells, two molecular scaffolds tether the KMN network to kinetochores: the constitutive inner kinetochore components CENP-T and CENP-C (Figure 1A) (Gascoigne et al. 2011, Przewloka et al. 2011, Nishino et al. 2013, Huis In ‘t Veld et al. 2016). CENP-C is recruited to centromeres by interacting with CENP-A nucleosomes via two nucleosome-binding domains (Carroll et al. 2010, Kato et al. 2013, Guo et al. 2017), whereas the C-terminus of CENP-T, in complex with CENP-W, -S, and –X, binds to the centromeric DNA (Hori et al. 2008, Nishino et al. 2012). The N-terminus of CENP-T can recruit one KMN complex via the Mis12-binding region; CENP-T N-tail can also recruit two Ndc80 complexes directly by binding to the Spc24/25 subunits (Huis In ‘t Veld et al. 2016). Recruitment of the Ndc80 and Mis12 complexes to CENP-T is promoted by phosphorylation of the CENP-T N-terminus by CDK1 mitotic kinase (Nishino et al. 2013, Rago et al. 2015, Huis In ‘t Veld et al. 2016). KMN complexes are also recruited to the kinetochore via the N-terminus of CENP-C, which has one Mis12-binding site (Petrovic et al. 2016), but the physiological significance of Ndc80 recruitment via different scaffolds in not fully understood (Schleiffer et al. 2012, Kim et al. 2015, Rago et al. 2015, Suzuki et al. 2015, Hara et al. 2018). The interaction between CENP-C and Mis12 complex is inhibited by the unphosphorylated N-terminal tail of Dsn1 subunit of Mis12 complex, which masks the CENP-C binding site on Mis12. This autoinhibition of the Mis12 complex is released when Aurora B kinase phosphorylates the Dsn1 tail (Yang et al. 2008, Kim et al. 2015, Rago et al. 2015, Petrovic et al. 2016).

The molecular and structural cues that orchestrate the hierarchical assembly of these components, and their specific contribution to microtubule binding, remain incompletely understood. In recent years, significant progress in dissecting kinetochore assembly has been achieved using artificial tethering of inner kinetochore components to Lac operator repeats integrated at non-centromeric regions in human and chicken cells (Barnhart et al. 2011, Gascoigne et al. 2011, Hori et al. 2013, Rago et al. 2015). Binding of fusions of LacI with proteins, such as the N-terminal fragments of CENP-T and CENP-C, or the CENP-A specific chaperone HJURP, leads to formation of ectopic kinetochore loci (reviewed in (Hori et al. 2020)). However, dissection of the underlying assembly pathways in living cells is difficult due to the presence of redundant interactions, which can complicate interpretation. A complementary strategy is to reconstruct kinetochore complexes using recombinant proteins, as demonstrated by the tour-de-force studies that assembled the entire linkage between human CENP-A nucleosomes, inner kinetochore components, and KMN (Figure 1B) (Weir et al. 2016, Pesenti et al. 2018). In a similar vein, a recent study using recombinant *Saccharomyces cerevisiae* proteins demonstrated that 14 CCAN subunits have the ability to self-assemble in the presence of CENP-A nucleosomes (Yan et al. 2019). Another study with yeast proteins revealed self-assembly of protein chains consisting of CENP-A nucleosomes, CENP-C and CENP-QU (Okp1/Ame1 complex) scaffolds, and the KMN components Mis12 and Ndc80 complexes (Hamilton et al. 2020). Such reconstituted chains bind to polymerizing microtubule ends and can withstand forces in 4–8 pN range (Hamilton et al. 2020). More complex microtubule end-coupling studies of these and the corresponding human complexes have lagged, in part due to difficulties with expression and purification of some kinetochore components. Another drawback is that recombinant components lack post-translational modifications that are known to regulate kinetochore assembly.

Kinetochore assembly has also been studied in cell extracts, which provide a rich source of kinetochore proteins with natural post-translational modifications (Figure 1B). In *Xenopus* egg extracts, soluble kinetochore components, including CENP-C and KMN, have been successfully recruited using sperm chromatin (Haase et al. 2017, Bonner et al. 2019) and CENP-A nucleosome arrays (Guse et al. 2011). The resultant assemblies exhibit microtubule binding and spindle checkpoint activities (Guse et al. 2011, Haase et al. 2017). Importantly, the critical regulatory step for kinetochore assembly in *Xenopus* egg extracts is N-terminal phosphorylation of the Dsn1 subunit of the Mis12 complex by Aurora B kinase (Haase et al. 2017, Bonner et al. 2019). Indeed, Mis12 complex containing phospho-mimetic substitutions in Dsn1 (S77E, S84E) allows robust assembly of functional kinetochores in *Xenopus* egg extracts even in the absence of Aurora B activity (Bonner et al. 2019). In yeast cell extracts, kinetochores can be assembled *de novo* using a 125-bp centromeric DNA as a template (Lang et al. 2018). In this system, recruitment of KMN components was increased by up to 7-fold using extracts from cells with phospho-mimetic mutations in Dsn1 (S240D, S250D). Interestingly, extracts prepared from yeast cells contain intact kinetochore particles that can be purified via a tag on Dsn1 (Akiyoshi et al. 2010, Gupta et al. 2018) (Figure 1B). These particles include KMN complexes and other outer kinetochore components, which enable microtubule binding (Akiyoshi et al. 2010). Load-bearing and microtubule tip tracking by these particles is significantly more robust than those of recombinant Ndc80 complexes alone (Powers et al. 2009, Akiyoshi et al. 2010), suggesting that these native particles are functionally active.

Although reconstructions in yeast and *Xenopus* egg extracts have been highly informative, there are significant differences in kinetochore organization and mitotic pathways between these systems and human cells. For example, in human cells, CENP-T is required for cell viability, whereas its yeast homologue, CNN1, is not (Bock et al. 2012). Also, protein scaffold Okp1/Ame1 is essential for cell viability and recruitment of KMN in yeasts (Hornung et al. 2014), but its human homologue CENP-QU is not (Bancroft et al. 2015, McKinley et al. 2015). Another important physiological difference is that the budding yeast kinetochore binds to a single microtubule (Winey et al. 1995), whereas human kinetochores have ∼20 microtubule binding sites (Wendell et al. 1993, McEwen et al. 2001), potentially imposing different requirements on assembly pathways. The *Xenopus* system appears to differ in the regulation of recruitment of Mis12 complex via the CENP-T pathway, which is controlled by Aurora B kinase in *Xenopus* cells (Bonner et al. 2019); in human cells, by contrast, recruitment is regulated by CDK1 kinase (Rago et al. 2015, Huis In ‘t Veld et al. 2016). Despite these and other differences, the success of reconstruction experiments in yeast and *Xenopus* extracts suggests that such reconstructions might also be possible in other mitotic systems. Hence, we adopted these strategies to examine kinetochore assembly pathways in extracts prepared from human cells.

## Results

### Kinetochore complexes in human mitotic cell extracts bind microtubules with low affinity

We first sought to investigate whether the cytosol of mitotic human cells contains stable native kinetochore particles that are capable of robust microtubule binding. To this end, HeLa cells stably expressing GFP fusions of individual components of Mis12 complex, Ndc80 complex, CENP-T, or CENP-C, were arrested in mitosis with nocodazole (Figure 1C). The arrested cells were harvested, lysed, and ruptured by gentle sonication. Next, the solution was clarified by centrifugation to obtain mitotic cell extracts. After dilution with lysis buffer, the concentrations of GFP-fused kinetochore proteins in the extracts, measured based on the intensity of GFP fluorescence, were in 10–100 nM range, reflecting the relative levels of these proteins in cells (Figure 1D).

We then tested whether these native GFP-labeled kinetochore complexes could interact with microtubules *in vitro*. To this end, we used a total internal reflection fluorescence (TIRF) microscopy– based assay to visualize binding of these complexes to fluorescently labeled microtubules stabilized with the non-hydrolyzable GTP analog GMPCPP (Figure 2A). Mitotic cell extracts from different cell lines were diluted in imaging buffer to obtain 3 nM GFP-fused proteins and then incubated with coverslip-immobilized microtubules (Figure 2B,C). Ndc80-GFP complexes exhibited frequent microtubule binding events, but they were not as robust as those observed for bacterially purified Bronsai Ndc80 complex (Supplementary Figure 1), which is a shortened Ndc80 complex with wild-type microtubule-binding domains (Wimbish et al. 2020). Microtubule binding by Bronsai Ndc80 was reduced when we added control cell extracts containing no GFP-fused proteins (Figure 2B,C; Supplementary Figure 1). However, even under these conditions, recombinant Ndc80 complex was still more active than native Ndc80 protein, suggesting that the difference in microtubule binding between these proteins was due to differences in their molecular structure or post-translational modifications, rather than the presence of other cytoplasmic proteins. Similarly poor binding was observed for native Ndc80 complexes in which GFP was fused to the Nuf2, Spc24, or Spc25 subunits, indicating that the partial inhibition was not caused by tag location.

**Figure 2.**
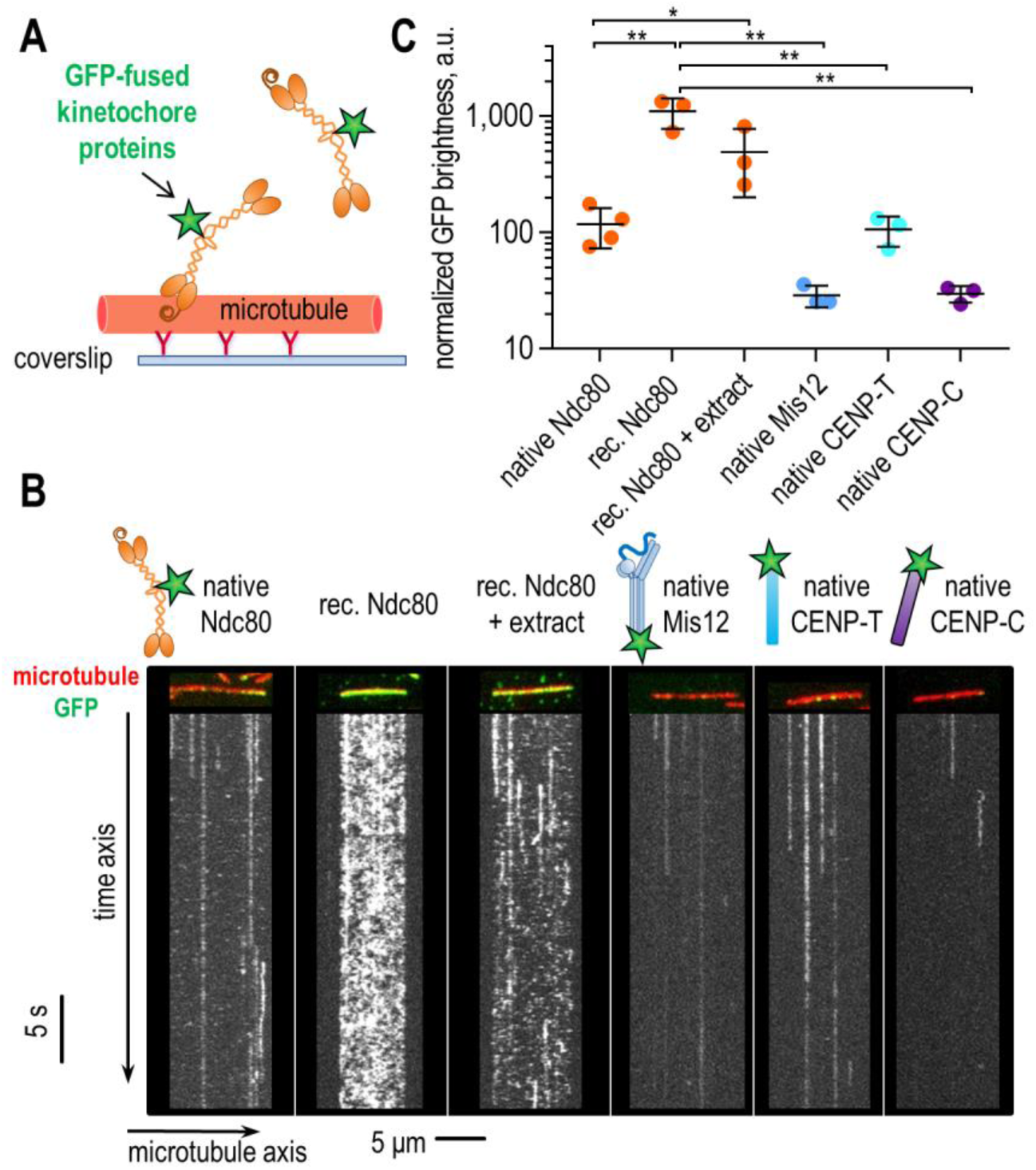
Native kinetochore complexes in the cytosolic fraction of mitotic cells bind microtubules only weakly. (A) Scheme of TIRF-based assay to visualize binding of soluble GFP-fused kinetochore components (2.5-3 nM) to GMPCPP-stabilized and fluorescently labeled microtubules, which were immobilized on the coverslip via anti-tubulin antibodies. (B) Colored images show representative microtubules (red) with bound GFP-fused kinetochore complexes (green); the corresponding gray-scale kymographs below reveal mobility of these complexes over 30-s observation time. (C) Average GFP brightness of microtubule decoration normalized against the concentration of GFP-labelled kinetochore protein (means with standard deviation (SD)); note semi-log scale; rec, recombinant. Each point represents an independent experiment in which brightness was collected from >18 microtubules. P-values were calculated by unpaired t-test: *, p<0.05; **, p<0.01. For more detailed statistics, see Source data.

We also used this assay to examine native GFP–labeled CENP-T, CENP-C, and Mis12 complexes. In these extracts, we detected some binding events between the GFP-labeled complexes and stabilized microtubules, but they were significantly rarer than with the native Ndc80 protein (Figure 2 B,C). Because CENP-T, CENP-C, and Mis12 complex do not have significant microtubule-binding activity on their own, these observations suggest that only a small fraction of these complexes is associated with microtubule-associated proteins, such as the Ndc80 complex. Alternatively, Ndc80 protein associated with these soluble GFP-labeled components might have decreased microtubule-binding activity.

### Native kinetochore complexes clustered on bead surfaces behave similarly to the soluble pool

Purified kinetochore particles from yeast readily bind microtubules whether they are soluble or conjugated to the surface of microbeads (Akiyoshi et al. 2010). To examine the activity of native human kinetochore complexes in a bead-bound context, we conjugated the complexes to 500-nm beads with anti-GFP antibodies (Figure 3A). These protein-coated beads were immobilized in microscopy chambers, washed to remove extracts, and incubated with fluorescently labeled stabilized microtubules in imaging buffer. As a positive control, we used beads coated with recombinant Bronsai Ndc80 complex, which readily recruited microtubules to the bead surface (Figure 3B), as expected (Powers et al. 2009, Chakraborty et al. 2019, Wimbish et al. 2020). As with soluble proteins, native Ndc80 complexes bound to the beads interacted with microtubules less strongly than Bronsai Ndc80 (Figure 3B-D). Thus, that bead binding does not improve Ndc80 activity, such as seen for Kinesin-1, which is autoinhibited in a soluble but not bead-bound form (Coy et al. 1999). Because recruitment of Ndc80 complex to Mis12 increases the affinity of Ndc80 for microtubules in the yeast system (Kudalkar et al. 2015, Scarborough et al. 2019), we expected that Mis12-GFP complexes conjugated to the bead surface would exhibit improved binding to microtubules relative to native Ndc80 complexes alone. However, in our assays with human extracts, beads coated with native GFP-labeled Mis12 complexes exhibited significantly reduced microtubule-binding than beads coated with native Ndc80 complexes. The poor microtubule binding by beads with conjugated human Mis12-GFP is in striking contrast to yeast system, where Mis12-containing complexes exhibit durable microtubule association (Akiyoshi et al. 2010). Human CENP-T and CENP-C complexes had similarly low microtubule-binding activity in our assay (Figure 3). Thus, individual kinetochore proteins and their preassembled complexes present in extracts prepared from mitotic HeLa cells have reduced functional activity.

**Figure 3.**
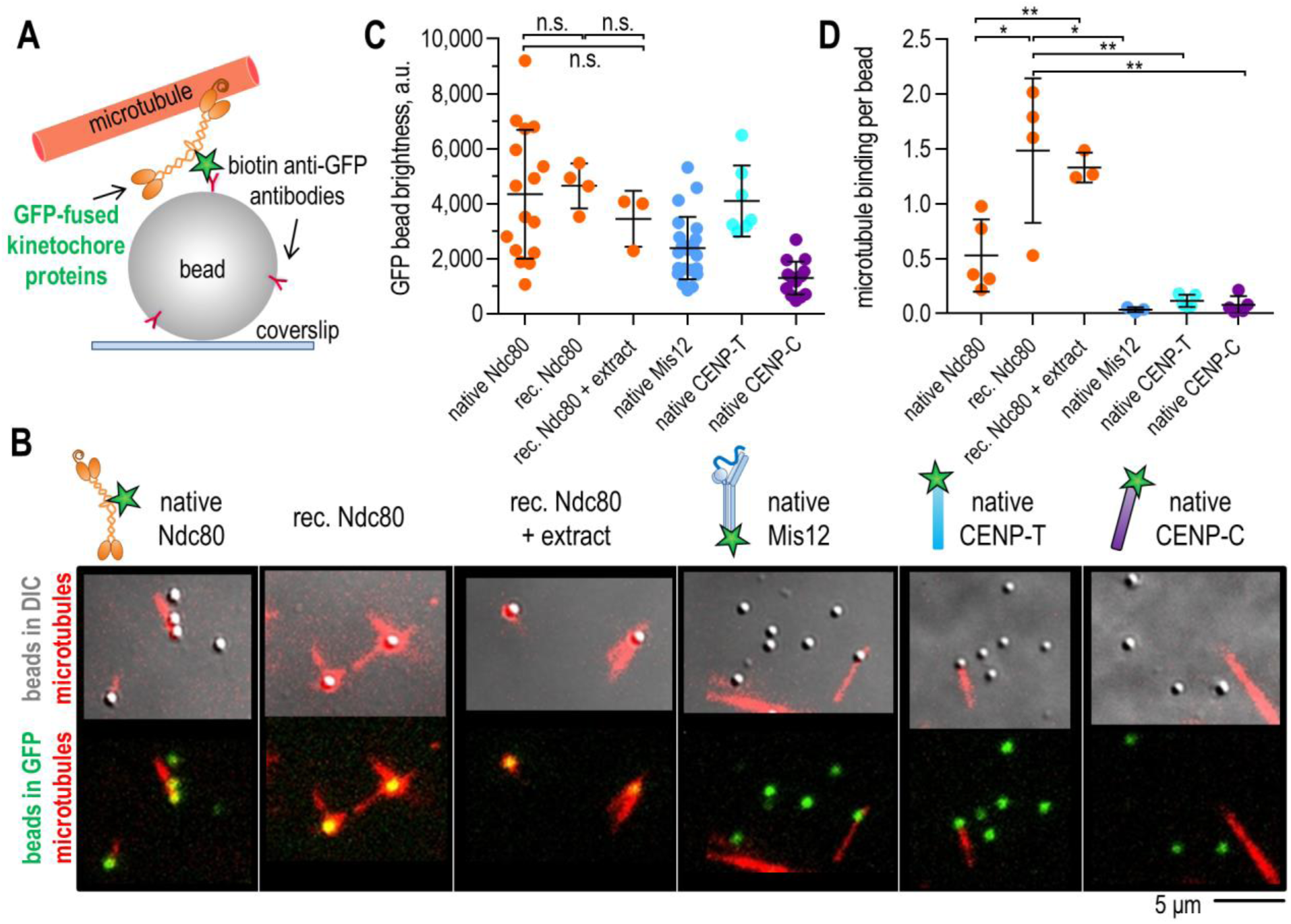
Clustering of native kinetochore complexes on microbeads does not enhance their binding to microtubules. (A) Scheme of bead-based assay to test microtubule binding of kinetochore components. (B) Representative examples of microscopy fields showing microtubules (red) and GFP-fused kinetochore components conjugated to coverslip-immobilized beads, shown in the DIC (top) and GFP (bottom) channels. (C) Average GFP bead brightness corresponding to the coating density of GFP-fused kinetochore components on bead surfaces. (D) Average number of bead-bound microtubules normalized to the number of beads per imaging field. On panels (C) and (D), means are shown with SD, and each point represents an independent experiment. P-values were calculated by unpaired t-test: n.s., p>0.05; *, p<0.05, **, p<0.01. For more detailed statistics, see Source data. Results for native Ndc80-GFP combine data from cell lines expressing GFP-fused Nuf2, Spc24, and Spc25.

### In mitotic cell extracts, recruitment of Ndc80 complexes via the CENP-T pathway is blocked

We next employed a different strategy in which we sought to assemble kinetochore particles *in vitro* using inner kinetochore components as nucleators for the assembly reactions. First, we purified full-length CENP-T/W complex, which can directly bind to the Ndc80 and Mis12 complexes (Gascoigne et al. 2011, Gascoigne et al. 2013, Nishino et al. 2013, Kim et al. 2015, Rago et al. 2015, Huis In ‘t Veld et al. 2016). To test the functionality of our recombinant construct, we immobilized CENP-T/W on beads via anti-CENP-T antibodies, and incubated these beads with recombinant Bonsai or Broccoli Ndc80 complexes that differed in their molecular compositions (Figure 4A). As expected, purified CENP-T/W complex recruited Bonsai Ndc80 complex (Nishino et al. 2013, Huis In ‘t Veld et al. 2016), which contains the kinetochore targeting domains from Spc24/25. By contrast, Broccoli Ndc80 complex, which lacks these domains (Schmidt et al. 2012), was not recruited (Figure 4B). Consistent with the importance of CENP-T phosphorylation for kinetochore assembly, binding of Bonsai Ndc80 complexes increased when we used a version of CENP-T/W complex containing three phospho-mimetic substitutions in the CDK target sites (T11D, T27D, and T85D), (Figure 4C) (Nishino et al. 2013, Huis In ‘t Veld et al. 2016, Hara et al. 2018).

**Figure 4.**
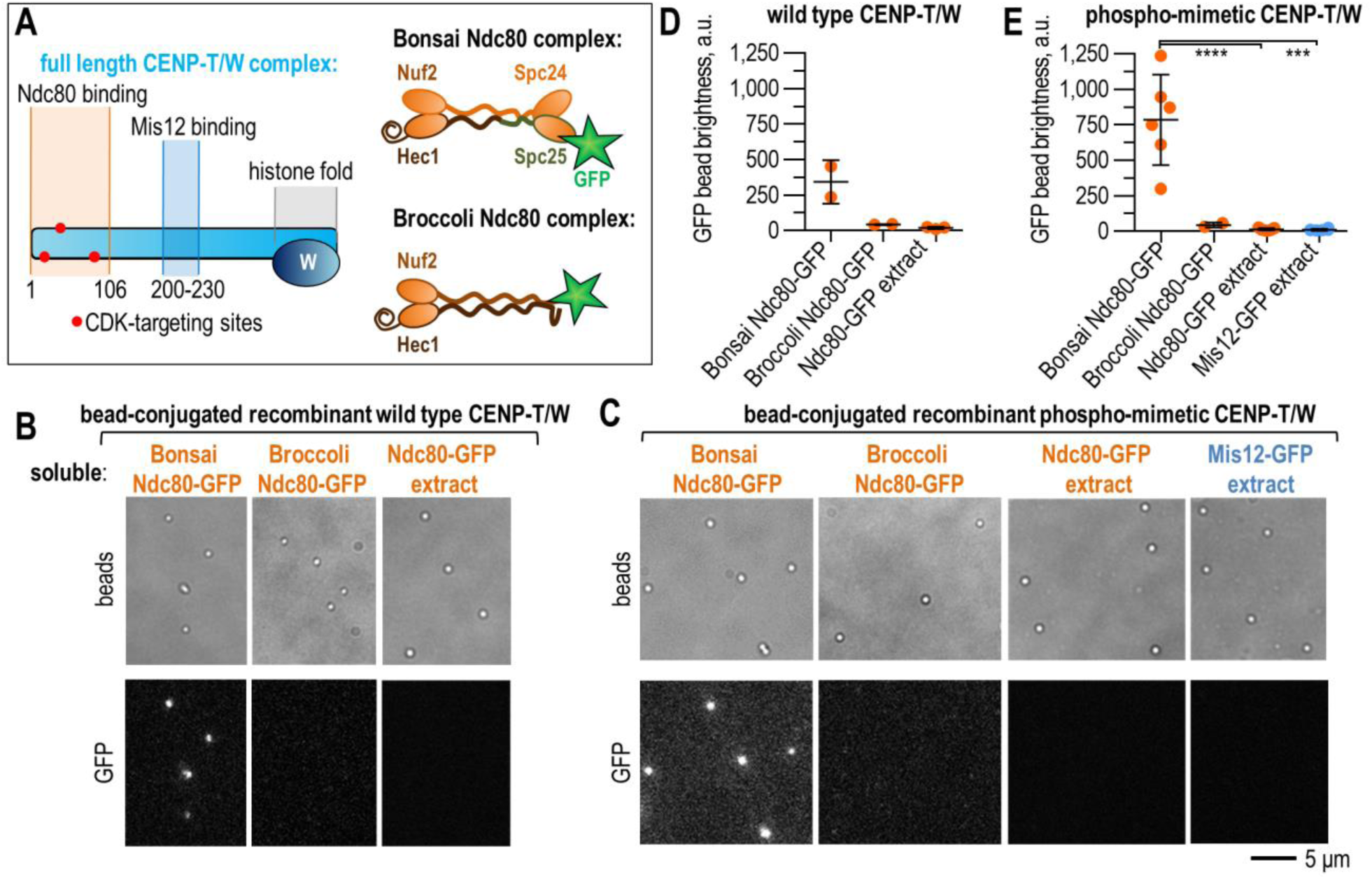
CENP-T robustly binds recombinant, but not native, Ndc80 complexes. (A) Schematic representation of recombinant CENP-T/W protein. The phospho-mimetic CENP-T/W construct contains substitutions, indicated by red dots (left). Schematic representation of recombinant Ndc80 constructs (right); see Supplementary Figure 1A for a comparison with wild-type Ndc80. (B and C) Representative images of coverslip-immobilized beads in bright field (top) and GFP (bottom) channels, showing recruitment of GFP-fused Ndc80 complexes to recombinant wild-type (B) or phospho-mimetic (C) CENP-T/W, which has no fluorescent tags. (D and E) Average GFP-brightness (mean with SD) of wild-type (D) or phospho-mimetic (E) CENP-T/W-coated beads incubated with various Ndc80 or Mis12 proteins, as in panels (B and C). Each point is derived from an independent experiment and represents the average brightness of >30 beads. P-values were calculated by unpaired t-test: ***, p<0.001; ****, p<0.0001. For more detailed statistics, see Source data. Concentrations of GFP-labeled soluble proteins, applied as minimally diluted mitotic cell extracts, were as follows: 120–130 nM recombinant Bonsai or Broccoli Ndc80, 50–190 nM native Ndc80, and 10-80 nM native Mis12 complexes.

Surprisingly, native GFP-fused Ndc80 complex, which was present in mitotic cell extracts at a concentration similar to that of the recombinant Ndc80 protein used in our assay, failed to bind CENP-T/W–coated beads. This lack of interaction was observed regardless of whether the beads were coated with wild type or phospho-mimetic CENP-T/W complex (Figure 4B-E), implying that CENP-T/W phosphorylation is not sufficient for this interaction. Recruitment of native Mis12-GFP to the CENP-T/W complex was also poor (Figure 4C,E), potentially due to the absence of phospho-mimetic substitutions in positions T195 and S201, which are important for Mis12-CENP-T binding in cells and *in vitro* (Rago et al. 2015, Huis In ‘t Veld et al. 2016). Together, these data establish that recombinant CENP-T/W complex with phospho-mimetic substitutions in three CDK-target sites (T11D, T27D, and T85D) cannot recruit soluble outer kinetochore components Ndc80 and Mis12 complexes. Because these substitutions are sufficient to promote robust binding between recombinant Ndc80 and CENP-T, we conclude that Ndc80 recruitment at the kinetochore requires additional cues that are absent in cell extracts.

### Recombinant CENP-A nucleosomes bind native CENP-C, but outer kinetochore complexes are not recruited

Next, we investigated whether functional complexes could be formed using centromere-specific CENP-A nucleosomes as templates for kinetochore assembly. We reconstituted recombinant CENP-A nucleosomes (Sekulic et al. 2016, Allu et al. 2019) using a Cy5-labled 147-bp sequence that corresponds to the preferred assembly site within a monomer of repetitive human centromeric α-satellite DNA (Hasson et al. 2013, Falk et al. 2015) (Figure 5A–C). Nucleosomes were immobilized at the surface of the antibody-coated beads via a His-tag engineered at the N-terminus of H2A histone (Figure 5D). Successful conjugation of these and similarly prepared H3 nucleosomes, which served as a negative control, was confirmed via fluorescence of the DNA-conjugated Cy5 fluorophore. Cell extract prepared from mitotic HeLa cells stably expressing GFP-fused CENP-C was incubated with the nucleosome-coated beads. GFP-CENP-C bound to beads with CENP-A nucleosomes (Figure 5E,F), consistent with the direct nature of this interaction (Carroll et al. 2010, Kato et al. 2013, Falk et al. 2015, Falk et al. 2016), but not with canonical H3 nucleosomes. The recruitment of CENP-C to CENP-A–coated beads is likely to be higher than what can be detected with GFP fluorescence, because both GFP-fused and untagged CENP-C proteins can bind to these nucleosomes. CENP-C enrichment on the beads was also observed using extracts prepared from DLD-1 cells in which both copies of the endogenous CENP-C gene were tagged with YFP (Supplementary Figure 2B), indicating that these interactions can take place at physiological levels of CENP-C. To determine the degree to which outer kinetochore components were recruited to this molecular platform, we incubated nucleosome-coated beads with extracts prepared from cells stably expressing GFP-fused Mis12 or Ndc80 complexes, in which CENP-C has no GFP tag. However, we observed little binding of Mis12 and Ndc80 complexes to the beads, and similarly low levels were seen for beads coated with CENP-A or H3 nucleosomes (Figure 5G, Supplementary Figure 2C,D). Thus, the CENP-A nucleosome–bound native CENP-C complex cannot facilitate recruitment of outer kinetochore components present in cell extracts.

**Figure 5.**
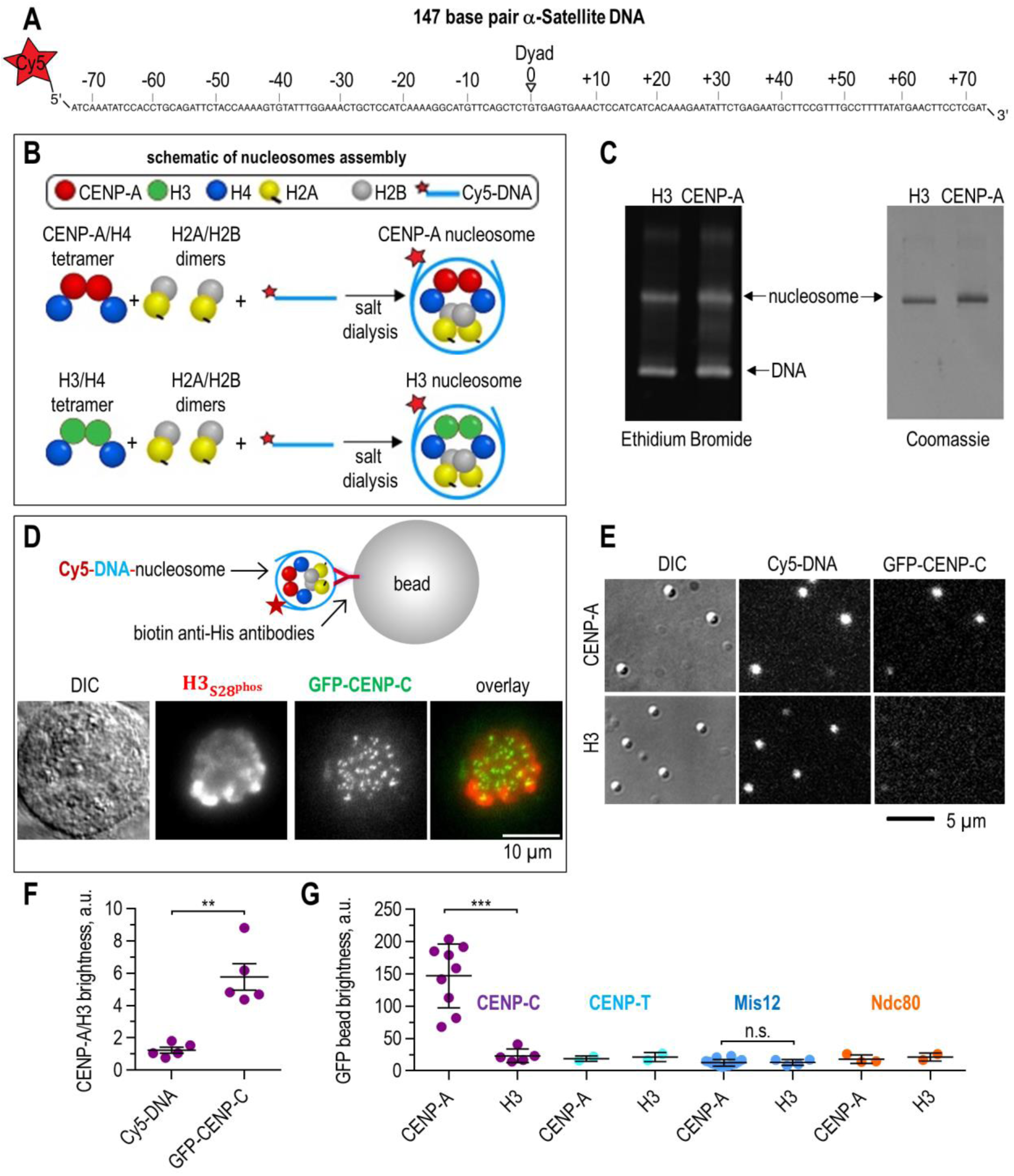
CENP-A nucleosomes specifically recruit native CENP-C, but not outer kinetochore components, from mitotic cell extracts. (A) The DNA (147 bp) used in nucleosome reconstitution is a CENP-A positioning fragment from the human X-chromosome centromeric α-satellite repeat; the 5’ end is labeled with Cy-5. (B) Schematic and constituents required to reconstitute nucleosomes with labelled DNA and His-H2A protein. (C) Nucleosomes prepared by gradient salt dialysis were analyzed by separating on 5% native PAGE gels followed by ethidium bromide and Coomassie staining. (D) Schematic of nucleosome immobilization on beads and an example of a mitotic HeLa cell stably expressing GFP-CENP-C, fixed and stained with anti-H3_S28_^phos^ antibodies to visualize chromosomes. (E) Representative images of beads in DIC and fluorescent channels, showing DNA-Cy5 and GFP-CENP-C following incubation with cell extracts. (F) Ratio of brightness of CENP-A– vs H3-coated beads in the Cy5 and GFP channels. The conjugation level of different nucleosomes is similar, but only nucleosomes containing CENP-A recruit native GFP-CENP-C. Means are shown with SEM. (G) Average GFP brightness of beads reflects the level of recruitment of the indicated kinetochore components to different nucleosomes. Means are shown with SD. Each point on panels (F and G) represents the average bead brightness obtained in one independent experiment. P-values were calculated by unpaired t-test: **, p<0.01; ***, p<0.001. For more detailed statistics, see Source data.

### Removal of the autoinhibitory domain of Mis12 complex partially improves its recruitment

To explore possible mechanisms preventing outer kinetochore assembly in cell extracts via the CENP-C pathway, we focused on the post-translational modifications of the Mis12 complex. Activation of this complex by phosphorylation of the N-terminal tail of Dsn1 is a key step for kinetochore assembly in *Xenopus* egg extracts (Bonner et al. 2019). We did not expect phosphorylation of the Dsn1 tail to be prevented in human mitotic cell extracts because they contain endogenous Aurora B kinase, and our lysis buffer included an ATP regeneration system. However, the activity of the soluble Aurora B kinase pool toward the Mis12 complex may be insufficient for robust Mis12 activation (Bonner et al. 2019). In *Xenopus, Saccharomyces cerevisiae, Kluyveromyces lactis*, chicken and human systems the requirement of Mis12 phosphorylation for CENP-C binding can be bypassed by introducing the truncations or phospho-mimetic substitutions at the Dsn1 N-terminus, thereby generating an autoinhibition-deficient Mis12 complex (Akiyoshi et al. 2013, Kim et al. 2015, Rago et al. 2015, Dimitrova et al. 2016, Petrovic et al. 2016, Hara et al. 2018, Lang et al. 2018, Bonner et al. 2019, Hamilton et al. 2020).

To explore the role of Mis12 phosphorylation in kinetochore assembly reactions in human cell extracts, we generated a HeLa cell line stably expressing GFP-fused Mis12 complex, in which the autoinhibitory region at the N-terminus of the Dsn1 subunit was deleted (GFP-Dsn1-Δ91-113) (Figure 6A). In previous studies (Kim et al. 2015) and in our experiments, this mutant exhibits constitutive kinetochore association in HeLa cells, indicating release of Mis12 autoinhibition (Supplementary Figure 3). Extracts prepared from mitotically arrested GFP-Dsn1-Δ91-113 cells were incubated with beads coated with either CENP-A or H3 nucleosomes (Figure 6B). Mutant complexes bound to these beads more strongly than wild-type Mis12-GFP complexes (Figure 6C). However, this increase could partially be attributed to a higher level of expression of mutant protein, which resulted in a 4-fold higher concentration of soluble phospho-mimetic Mis12 relative to wild-type complex (Supplementary Figure 4D). Therefore, while relieving autoinhibition of Mis12 complex improves its recruitment to kinetochores, phosphoregulation of the Dsn1 tail is not the major factor limiting binding of the Mis12 complex to the CENP-A nucleosome–bound CENP-C in human cell extracts.

**Figure 6.**
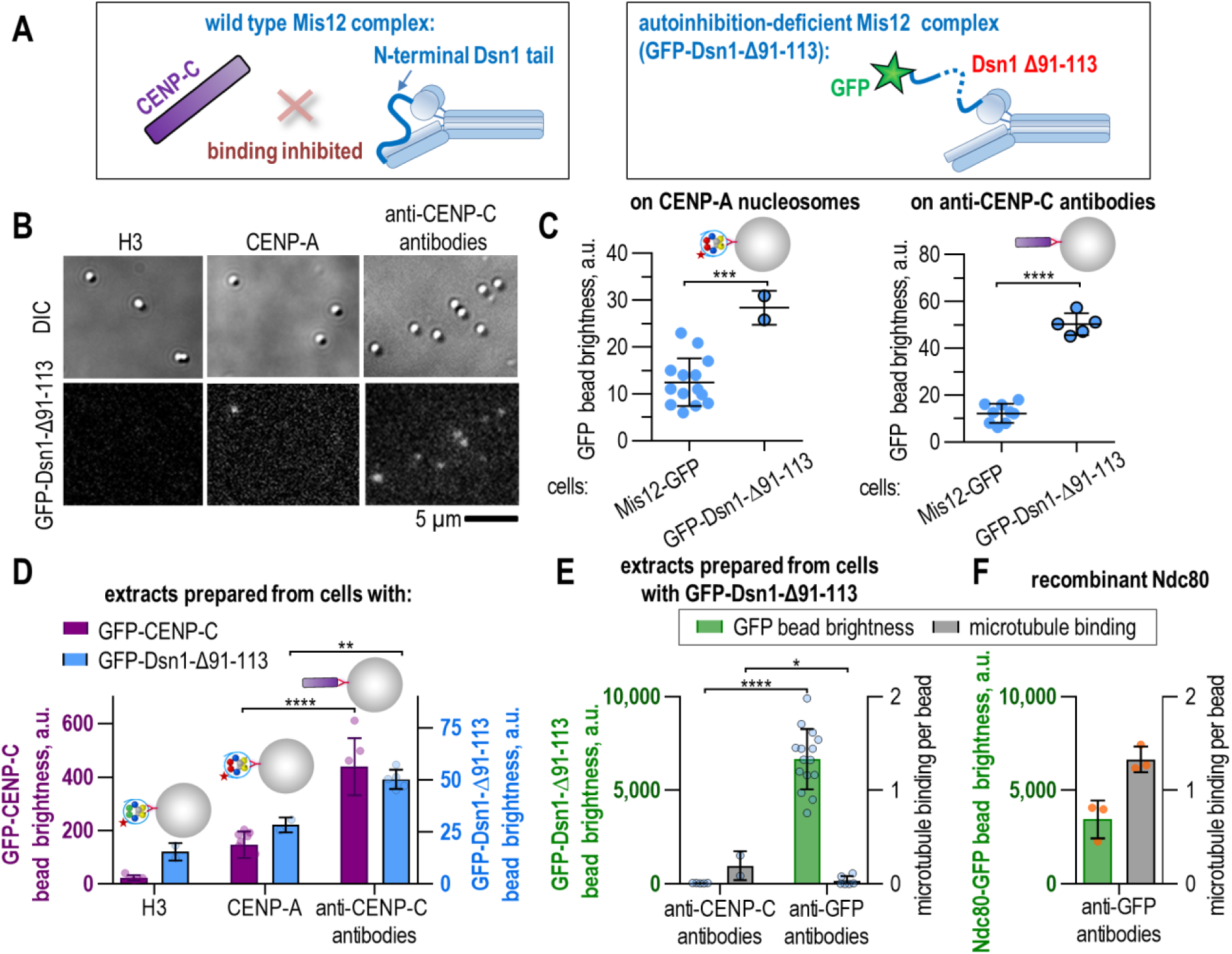
Testing of the autoinhibition-deficient Mis12 complex. (A) Cartoon illustrating autoinhibition of the Mis12 complex and its activation by truncation of a segment of the Dsn1 N-tail. (B) Representative images of beads in the DIC and GFP channels, showing recruitment of native GFP-fused Dsn1-Δ91-113 to H3- and CENP-A nucleosomes, or to beads coated with anti–CENP-C antibodies. (C) GFP bead brightness showing recruitment of native GFP-fused Mis12 or Dsn1-Δ91-113 to beads via CENP-A nucleosomes (left) or anti–CENP-C antibodies (right). (D) Average GFP-brightness of beads showing recruitment of native GFP-fused CENP-C (left axis) and Dsn1-Δ91-113 (right axis) to beads with indicated coatings; data for Dsn1-Δ91-113 is the same as on panel (C), plotted here to provide side-by-side comparison with the recruitment of native CENP-C. (E) Average GFP-bead brightness (left axis and green columns) and the number of bead-bound microtubules (right axis and clear columns) plotted for beads coated with GFP-Dsn1-Δ91-113 (blue dots) directly via anti-GFP antibodies or indirectly via anti–CENP-C antibodies. Data regarding GFP brightness of beads coated with GFP-Dsn1-Δ91-113 via anti-CENP-C antibodies are the same as in panel (C). (F) Results of experiments similar to those in panel (E), but using recombinant Ndc80 (orange dots) in the presence of unlabeled cell extract. These data are the same as in Figure 3C,D, and are shown here for side-by-side comparison with the recruitment activity of native kinetochore complexes. On panels (C-F), horizontal lines and column bars show means with SD, each dot represents the average result from one independent experiment. P-values were calculated by unpaired t-test: *, p<0.05, **, p<0.01; *** p<0.001; p ****, p<0.0001. For more detailed statistics, see Source data.

To further test this conclusion, we used a different Mis12 mutant in which two phospho-mimetic substitutions (S100D and S109D) were introduced into the N-terminal tail of the Dsn1 subunit in DLD-1 cells (Supplementary Figure 5). Phospho-mimetic Dsn1 fused to GFP (GFP-Dsn1-SD) was introduced at a unique genomic locus using Flp/FRT recombination. GFP-Dsn1-SD constitutively localized to kinetochores during interphase and mitosis, consistent with previous findings in human cells (Gascoigne et al. 2013, Kim et al. 2015, Rago et al. 2015). However, the recruitment of these native phospho-mimetic Mis12 complexes to CENP-A nucleosomes was poor, whereas the binding of native CENP-C from DLD-1 extracts to CENP-A nucleosomes was readily detected (Supplementary Figure 6). The relative inefficiency of the phospho-mimetic mutations in our assay is consistent with previous observations that phospho-mimetic Dsn1 mutant has weaker CENP-C binding than that see with N-terminal truncations in Dsn1 (Dimitrova et al. 2016, Petrovic et al. 2016).

### Neither elevated density of bead-bound CENP-C nor the combination of CENP-C and CENP-T scaffolds are sufficient for robust recruitment of outer kinetochore proteins

Recruitment of native Mis12 complexes to beads containing CENP-C bound to CENP-A nucleosomes could be limited by the relatively low surface density of CENP-C protein on these beads, obscuring the recruitment of Mis12 complex in our experimental system. To test this idea, we coated beads with antibodies against CENP-C, and then incubated them with extracts from mitotic HeLa cells expressing GFP-CENP-C. The GFP bead brightness increased about 2-fold relative to beads that recruited GFP-CENP-C via CENP-A nucleosomes (Figure 6D). We then incubated the antibody-coated beads with extracts containing either Mis12-GFP or GFP-Dsn1-Δ91-113 to monitor the recruitment of the GFP fusion proteins to these beads via the unlabeled CENP-C present in these extracts (Figure 6B). In agreement with proposed autoinhibition of the Mis12 complex, we observed increased recruitment of GFP-Dsn1-Δ91-113, but not wild-type Mis12-GFP, to these beads (Figure 6C right graph).

Next, we investigated whether this improved recruitment of truncated Mis12 complex could enhance the downstream assembly reactions. To assess recruitment of the outer kinetochore proteins with microtubule-binding activity to the autoinhibition-deficient Mis12 complex, we first used beads coated with anti-CENP-C. After incubation with the GFP-Dsn1-Δ91-113 cell extracts, we incubated the beads with stabilized fluorescent microtubules and scored the number of microtubules bound to the beads, as depicted in Figure 3. Microtubule binding by these beads, as well as those coated with anti-GFP antibodies, was poor relative to the positive control, which used beads coated with recombinant Ndc80 complexes (Figure 6 E,F; Supplementary Figure 4C). Second, we investigated whether soluble GFP-Dsn1-Δ91-113 protein could form complexes with active microtubule-binding proteins. For this purpose, we incubated extracts containing GFP-Dsn1-Δ91-113 with stabilized microtubules and imaged via TIRF microscopy. Soluble GFP-fused Dsn1-Δ91-113 complexes behaved similarly to wild-type Mis12-GFP, showing relatively weak affinity for microtubules (Supplementary Figure 4A,B). These findings suggest that the autoinhibition of Mis12 complex by N-terminal tail of Dsn1 has no effect on the recruitment of Ndc80 complex, in agreement with findings in yeast (Akiyoshi et al. 2013). Importantly, native GFP-Dsn1-Δ91-113 complexes in the soluble pool behaved similarly to complexes clustered on beads via the CENP-C scaffold or anti-GFP antibodies, indicating that their inability to recruit active outer kinetochore components in cell extracts is a property of these complexes, rather than the result of the specific mode of their recruitment to the beads.

Finally, we investigated whether kinetochore assembly in cell extracts can be activated by a combination of components from different recruitment pathways. To create bead coatings with more complex molecular compositions, we incubated beads with a mixture of anti-CENP-C and anti-CENP-T antibodies, and then incubated them with extracts prepared from mitotically arrested HeLa cell lines expressing GFP-Dsn1-Δ91-113. However, this approach, did not appreciably improve recruitment of GFP-Dsn1-Δ91-113 (Supplementary Figure 4E). We observed similarly low recruitment when we coated beads with a mixture of CENP-A and H3 nucleosomes. Consistent with this, beads with more complex molecular coatings bound microtubules as rarely as beads coated with only one inner kinetochore component (Supplementary Figure 4F). Thus, combining different molecular nucleators is not sufficient to overcome the multiple inhibitory steps that limit spontaneous kinetochore assembly in mitotic cell extracts.

## Discussion

Inspired by previous kinetochore reconstruction studies in yeast and frog egg systems (Akiyoshi et al. 2010, Guse et al. 2011, Haase et al. 2017, Lang et al. 2018, Bonner et al. 2019, Hamilton et al. 2020), we applied analogous strategies to investigate the assembly and microtubule-binding activities of the cytoplasmic pool of kinetochore proteins in mitotically arrested human cells (Figure 7A). Native GFP-fused kinetochore proteins, such as the Ndc80 and Mis12 complexes and the CENP-T and CENP-C scaffold, were present at concentrations of 10–100 nM in the soluble fraction of extracts prepared from HeLa cells stably expressing these proteins. Using multiple microscopy-based assays *in vitro*, we investigated the ability of these components to form active kinetochore protein assemblies *de novo*. Surprisingly, we found that only a few kinetochore assembly steps were permitted in human mitotic cell extracts, whereas many binding reactions were either inefficient or blocked completely.

**Figure 7.**
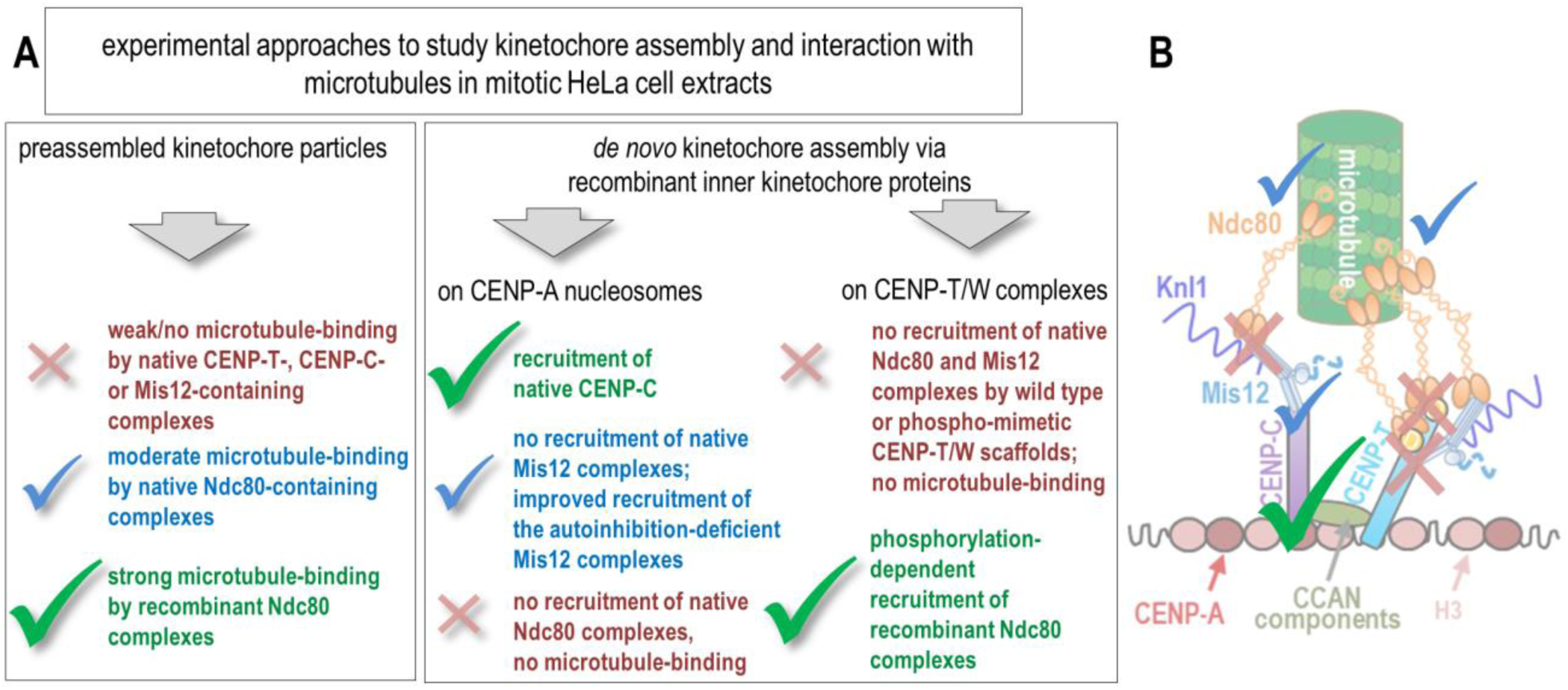
Summary of the permitted and restricted assembly steps in human cell extracts. (A) Experimental approaches to reconstitute kinetochore assembly and function *in vitro* using mitotic HeLa cell extracts and the main conclusions derived from this study. (B) Simplified scheme indicating the permitted assembly steps (green check-mark), moderately/weakly efficient reactions (blue check-mark), and restricted interactions (red crosses). Symbols “D” indicate phospho-mimetic substitutions in CENP-T.

The CENP-C–dependent pathway can be partially reconstructed in human cell extracts: recombinant CENP-A nucleosomes conjugated to the bead surface, but not canonical H3-containing nucleosomes, recruit native CENP-C (Figure 5 E,F), consistent with a direct interaction between recombinant CENP-A and CENP-C (Carroll et al. 2010, Kato et al. 2013, Falk et al. 2015, Falk et al. 2016). However, recruitment of the outer kinetochore components Mis12 and Ndc80 complexes to this molecular platform is poor. These *in vitro* findings are in agreement with the results of kinetochore reconstruction studies in HeLa cells, in which overexpression of CENP-A promotes its incorporation throughout the chromatin and subsequent recruitment of inner kinetochore proteins (CENP-C, CENP-N and Mis18), but not the KMN components (Gascoigne et al. 2011). By contrast, the KMN components can be recruited to single non-centromeric CENP-A loci generated by tethering a fusion of LacI with the HJURP histone chaperone to LacO arrays (Barnhart et al. 2011, Logsdon et al. 2015). It is currently not known why the efficiency of outer kinetochore assembly differs among these various cellular approaches, and whether similar mechanisms can explain the *in vitro* results. Failure of KMN recruitment following CENP-A overexpression could result from the limited size of soluble pools of KMN components, preventing their visible accumulation on the chromosome arms. As explained below, however, in our reconstructions, the size of soluble protein pools was not a limiting factor. An alternative explanation for the different extents of KMN recruitment in various cellular systems is that proper kinetochore assembly and maturation rely on some specific cues provided by the structural and molecular environment of CENP-A-containing chromatin at the sites with a locally high density of CENP-A nucleosomes. In this case, the failure of bead-bound CENP-A nucleosomes to nucleate robust assembly of soluble kinetochore components in mitotic cell extracts could be explained by the lack of a proper milieu or some crucial regulatory modifications. Future work is needed to test these ideas, and to identify and recapitulate the missing factors or regulatory steps.

Work in other systems identified phosphorylation of Dsn1 by Aurora B kinase as a key step that partially or completely relieves Mis12 autoinhibition, promoting its binding to kinetochore-localized CENP-C, followed by recruitment of Ndc80 complex and other outer kinetochore proteins (Yang et al. 2008, Kim et al. 2015, Rago et al. 2015, Petrovic et al. 2016, Lang et al. 2018, Bonner et al. 2019). Indeed, when we used mitotic cell extracts containing autoinhibition-deficient Mis12 complex (GFP-Dsn1-Δ91-113), which localizes constitutively to HeLa kinetochores, binding of Mis12 complexes to the CENP-A–coated beads was noticeably improved relative to wild-type Mis12. However, recruitment was still disappointingly inefficient, as only one mutant Mis12 complex was recruited per six native CENP-C molecules associated with the bead-bound CENP-A nucleosomes (Figure 6D). The total level of GFP-Dsn1-Δ91-113 binding to beads was improved by increasing the density of bead-bound CENP-C molecules, when conjugations were carried out using anti-CENP-C antibodies. When native GFP-CENP-C was tethered directly to the beads with anti-CENP-C antibodies, the level of GFP-CENP-C bound to the beads was 2-fold higher than what we could achieve with recombinant CENP-A nucleosomes. However, even in this case, there was only one autoinhibition-deficient Mis12 complex per nine CENP-C molecules (Figure 6D), strongly suggesting that this interaction is not limited by the level of bead-bound CENP-C. It is also not limited by the availability of soluble Mis12 complexes, as recruitment of wild-type or mutant Mis12 complexes to beads directly via anti-GFP antibodies led to a >15-fold increase in bead brightness relative to Mis12 recruitment via native CENP-C. These results strongly imply that phosphorylation of Dsn1 is not the main factor that limits interaction between the native Mis12 complex and CENP-C proteins in human cell extract.

Interestingly, autoinhibition-deficient Mis12 complexes were unable to promote recruitment of native Ndc80 complexes, as seen from the low microtubule binding activity of beads coated with Mis12 complexes via different means: via anti-GFP antibodies, via the CENP-C bound to beads directly (with anti– CENP-C antibodies), or via bead-bound CENP-A nucleosomes (Figure 6E,F). Consistent with this, only a minor fraction of soluble Mis12-containing complexes in mitotic cell extracts have microtubule-binding activity (Figure 2B,C; Supplementary Figure 4A,B). Together, these results strongly suggest that extracts prepared from mitotic HeLa cells do not contain significant amounts of Mis12-containing preassembled kinetochore particles, such as those found in yeast cell extracts.

Also unexpected was the lack of robust interactions between recombinant CENP-T and native Ndc80 complexes. Studies in different organisms found that CDK1 phosphorylation in the Ndc80-binding region of CENP-T is required for recruitment of two Ndc80 complexes per CENP-T (Gascoigne et al. 2011, Nishino et al. 2013, Rago et al. 2015, Huis In ‘t Veld et al. 2016, Hara et al. 2018). Weak binding between native Ndc80 complex and recombinant full-length CENP-T/W complex containing key phospho-mimetic substitutions (T11D, T27D, T85D) is unlikely to be explained by the lack of natural phosphoryl groups, as this mutant CENP-T construct readily recruits recombinant Ndc80 protein containing Spc24/25 domains (Figure 5E,F). Robust recruitment of Ndc80 complex to kinetochores in mitotic cells could therefore require additional post-translational modifications at sites not targeted in our mutant, or the release of autoinhibition of native Ndc80 complex by a mechanism that remains to be elucidated.

In summary, using different approaches, we found that multiple inhibitory steps prevent outer kinetochore assembly in mitotic cell extracts (Figure 7B). At this time, we cannot exclude the possibility that our experimental conditions are not a sufficiently close match to the molecular milieu for kinetochore assembly reactions in live cells. However, because consistent results were obtained with different approaches and for several kinetochore components, our findings strongly suggest that the absence of active preassembled kinetochore complexes and their *de novo* formation in cell extracts reflect the presence of physiological barriers to the assembly reactions involving the cytosolic pool of kinetochore components. We hypothesize that such inhibitory mechanisms guard against spurious formation of kinetochore-like structures in cell cytoplasm, which would compete with the formation of endogenous kinetochores and interfere with chromosome motility, spindle function, or checkpoint signaling.

The success of analogous reconstructions in other organisms could be explained by their different mitotic physiology, which appears to be advantageous for reconstitution assays, but also represents a source of variations in regulation of kinetochore assembly pathways. Although functional kinetochore particles containing the majority of known kinetochore proteins can be purified from yeast extracts (Gupta et al. 2018), the exact origin of these macromolecular assemblies remains unclear. They could represent a nucleoplasmic pool of kinetochore subcomplexes, or become extracted from exogenous centromeric location during cell lysis. In our experiments, kinetochore proteins could be extracted from the pool of DNA-bound proteins, as ruptured cells were treated with endonuclease to digest chromosomal DNA. It is more likely, however, that these proteins represent the pool of soluble cytoplasmic kinetochore components that are readily detected in the cytoplasm of dividing cells via the GFP fusions. The brightness of GFP-labeled kinetochore complexes visualized on microtubules *in vitro* suggests that they contain just one or a few GFP fluorophores, consistent with the idea that our findings represent properties of the cytoplasmic pool of kinetochore complexes, rather than fragments of human kinetochores. By contrast, kinetochore particles that spontaneously assemble in yeast cells contain multiple copies of the majority of kinetochore proteins (Gonen et al. 2012). We hypothesize that nucleation of yeast kinetochores at a defined short DNA sequence, as opposed to the megabases of repetitive DNA in human cells, could render yeast cells more tolerant to the presence of active kinetochore subcomplexes; this issue deserves further investigation. Also, the persistent attachment of assembled yeast kinetochore to microtubules throughout the cell cycle suggests that their incorporation into new kinetochores is regulated by different mechanisms than in human cells.

*Xenopus* egg extracts are also quite different from somatic human cells, as they contain concentrations of kinetochore proteins sufficient to assemble many thousands of kinetochores. In this system, the rate-limiting step is phosphorylation of the Dsn1 subunit of Mis12 complex, which takes place only at kinetochores due to the coincidence of CENP-C and high local concentrations of Aurora B kinase, which stabilizes the otherwise transient Mis12-CENP-C interaction (Bonner et al. 2019). Similar *in situ* reactions may be required to activate other loosely associated kinetochore components, thereby permitting their strong recruitment by maturing human kinetochores. Successful assembly of human kinetochores *de novo* from native proteins will require reconstruction of these as-yet-unknown regulatory steps and molecular environments, paving the way to a deeper mechanistic understanding of kinetochore formation in human cells.

## Materials and Methods

### Cell lines

HeLa and DLD-1 cells were cultured in Dulbecco’s modified Eagle’s medium with 10% fetal bovine serum and 1% penicillin–streptomycin at 37°C in a humidified atmosphere with 5% CO_2_. HeLa cells stably expressing GFP-fused kinetochore proteins (Mis12, CENP-T, Spc25, CENP-C, Nuf2 and Spc24) were kindly provided by the Lampson lab (University of Pennsylvania). The Mis12-GFP-Halo cell line was created using CRISPR-Cas9 to tag the endogenous Mis12 with GFP, as described in (Zhang et al. 2017). Halo-GFP-CENP-T and Halo-GFP-Spc25 cell lines were created using the recombinase-mediated cassette exchange system (Zhang et al. 2017). Analogous procedures were used to generate HeLa cell lines expressing Halo-GFP-CENP-C, Nuf2-GFP-3xHalo, and 3xHalo-GFP-Spc24. Briefly, HeLa acceptor cells were grown in 6-well plates to 60–80% confluence. Cells were co-transfected with 1 μg of the corresponding plasmids and 10 ng of a Cre recombinase plasmid using Lipofectamine 2000 (Thermo Fisher Scientific). Stable cell lines were selected after 2 days of growth by addition of 1 μg ml^−1^ puromycin. The unlabeled cell line used in this study was the HiLo acceptor HeLa cell line (Zhang et al. 2018). Additionally, we generated a clonal cell line stably expressing Mis12 complex with an N-terminal GFP-tag and truncated Dsn1 subunit (Δ91–113). This line was generated by retroviral infection of HeLa cells with a pBABE-blast–based vector, similarly to the procedure described in (Gascoigne et al. 2013). DLD-1 Flp-In T-REx cells stably expressing CENP-C^AID-EYFP/AID-EYFP^ was generated as described in (Fachinetti et al. 2015). To create the DLD-1 cell line expressing GFP-Dsn1 with two phospho-mimetic substitutions (Dsn1-SD), we generated a plasmid encoding human Dsn1 with the S100D and S109D mutations. This plasmid and FLP recombinase were transfected into DLD-Flp-In-T-REx cells (Supplementary Figure 5A). Hygromycin selection was performed for 10 days, followed by analysis of Dsn1-SD localization in hygromycin-resistant colonies. Kinetochore localization of GFP-Dsn1 was confirmed by fluorescence microscopy (Supplementary Figure 5B). GFP-fused proteins in other cell lines also exhibited prominent kinetochore localization.

### Immunofluorescence of cultured cells

To verify kinetochore localization of tagged proteins in HeLa cells, the cells were plated on glass coverslips coated with poly-D-lysine (Electron Microscopy Sciences). To enrich mitotic cells, cells were treated with 100 ng ml^−1^ nocodazole for 14 h before fixation. Cells were fixed with 3.7% paraformaldehyde (Electron Microscopy Sciences) in Corning Dulbecco’s phosphate-buffered saline without calcium and magnesium (Corning PBS) for 10 min. Next, cells were treated with 0.1% Tween-20 (Sigma-Aldrich) for 10 min. For DNA staining, mouse anti-H3_S28_^phos^ antibodies from Santa Cruz Biotechnology were used at 1:200 dilution in Corning PBS supplemented with 4 mg ml^−1^ bovine serum albumin (BSA; Sigma-Aldrich). Cell samples were washed and blocked with Corning PBS containing 10 mg ml^−1^ BSA and 1.5 mg ml^−1^ casein (Sigma-Aldrich, C5890) for 30 min. Secondary antibodies were Alexa Fluor 647–conjugated anti-mouse (Abcam) used at a dilution of 1:100. Samples were incubated with antibodies for 1 h at room temperature, and primary and secondary antibodies were ultracentrifuged (157,000 *g*, 15 min, 4°C) before application to remove aggregates. For Figure 1D, DNA staining with 500 nM propidium iodide (Molecular probes) was carried out after treatment with 50 μg ml^−1^ RNase (Qiagen) for 20 min at 37°C. All samples were mounted in VECTASHIELD (Vector Laboratories). For Supplementary Figure 3, DNA was stained with 4′,6-diamidino-2-phenylindole (DAPI, Sigma-Aldrich). Imaging of cells was carried out in epifluorescence and differential interference contrast (DIC) modes using a Nikon Eclipse Ti-E inverted microscope equipped with a 1.49 NA 100× oil objective and Andor iXon3 EMCCD camera, as described in (Chakraborty et al. 2018). Z-stacks were taken with 0.5 μm spacing for a total of 5 μm in the DIC, GFP, and DNA channels. All cell images are maximum-intensity projections from z-stacks, prepared using the Fiji software (Schindelin et al. 2012).

For Dsn1-SD localization studies in DLD-1 cells, immunofluorescence was performed on either asynchronous or mitotically arrested cells, which were obtained by adding 50 μM S-Trityl-L-cysteine for 14 h before fixation. Cells were fixed with 4% formaldehyde (Tousimis) for 10 min, quenched with 100 mM Tris (pH 7.5), and permeabilized with 0.5% Triton X-100 in Corning PBS for 5 min. Coverslips were washed three times in 0.1% Tween-20 in Corning PBS and incubated with blocking buffer (Corning PBS containing 2% fetal bovine serum, 2% BSA, and 0.1% Tween-20) for 5 min prior to primary antibody incubations. Incubation with anti-centromere antibody (1 μg/mL; Antibodies Inc. #155-235) was performed at room temperature in blocking buffer for 1 h, followed by incubation with Cy3-conjugated anti-rabbit secondary antibody (Jackson ImmunoResearch Laboratories #111-165-144) at a dilution of 1:200. DNA was stained with DAPI, and coverslips were mounted in VECTASHIELD (Vector Laboratories). Imaging of cells was captured on an inverted fluorescence microscope (DMI6000 B; Leica) at 100x, 1.4 NA oil immersion objective, and set with a charge-coupled device camera (ORCA AG; Hamamatsu Photonics). Z-stacks were collected at 0.2-μm spacing for a total of 6 μm in the GFP, Cy3, and DAPI channels. Cells were cropped, z-stacks were deconvolved using the LAS-AF software (Leica), and images were prepared using the Fiji software.

### Preparation of mitotic cell extracts

Cells were arrested with 100 ng ml^−1^ nocodazole for 14 h, at which time ∼60% of cells were mitotic, as judged by DIC and DNA visualization. Mitotic cells were harvested by shaking off and gentle rinsing of 15-cm tissue culture plates using a pipette. Harvested cells were pelleted and washed by centrifugations at 1,000 *g* in Corning PBS buffer. Cells were resuspended in 50 mM HEPES pH 7.2, 2 mM MgCl_2_, 150 mM K-glutamate, 0.1 mM EDTA, 2 mM EGTA, 10% glycerol, and washed by centrifugation; cell pellets containing 0.2-0.4×10^8^ cells were snap-frozen and stored in liquid nitrogen. Prior to each experiment, one cell pellet (volume 150–300 µl) was resuspended in two volumes of ice-cold lysis buffer (50 mM HEPES pH 7.2, 2 mM MgCl_2_, 150 mM K-glutamate, 0.1 mM EDTA, 2 mM EGTA, 0.1% IGEPAL, 10% glycerol, 4 mM Mg-ATP, 2 mM DTT) supplemented with protease inhibitors (0.2 mM 4-(2-Aminoethyl)-benzenesulfonylfluoride hydrochloride (Goldbio), 10 μg ml^−1^ leupeptin (Roche), 10 μg ml^−1^ pepstatin (Roche), 10 μg ml^−1^ chymostatin (Sigma-Aldrich), Complete Mini EDTA free cocktail (Roche)), phosphatase inhibitors (100 ng ml^−1^ microcystin-LR (Enzo Life Sciences), 1 mM sodium pyrophosphate (Sigma-Aldrich), 2 mM sodium-beta-glycerophosphate (Santa Cruz Biotechnology), 100 nM sodium orthovanadate (Alfa Aesar), 5 mM sodium fluoride (Sigma-Aldrich), 120 nM okadaic acid (EMD Millipore), PhosSTOP cocktail (Roche))], ATP regeneration system (10 mM creatine, 0.45 mg ml^−1^ phospho-creatine kinase (Sigma-Aldrich). During initial stages of this work, extracts were prepared using modified lysis buffer with lower ionic strength and detergent concentration (50 mM HEPES, pH 7.2,2 mM MgCl_2_, 0.1 mM EDTA, 2 mM EGTA, 0.05% IGEPAL, 10% glycerol) supplemented with Complete Mini EDTA free cocktail and PhosSTOP cocktail. In efforts to improve activity of cell extracts, we also varied the concentration of Mg-ATP and omitted ATP regeneration system and endonuclease. None of these conditions affected the recruitment of CENP-C, Mis12, or Ndc80 complexes to nucleosomes. In addition, there was no significant effect of these modifications on reconstitution experiments employing recombinant CENP-T/W, anti-CENP-C or anti-GFP antibodies, so data from experiments using different buffer compositions were combined. In several experiments, cells arrested in nocodazole were also treated for 1 h before harvest with 2.5 µM ZM447439, an Aurora B inhibitor (Tocris). The inhibitor had no significant effect on the levels of CENP-C, Nuf2, or Mis12 recruitment to CENP-A nucleosomes in cell extracts (Supplementary Figure 2A). Accordingly, the data from these two conditions were combined.

For all samples, cell pellets in lysis buffer were melted on ice for 5–10 min and homogenized by pipetting. Immediately thereafter, the cells were ruptured by sonication using a Branson SFX150 Sonifier with a 3/32” microtip and following settings: 68% power for five cycles consisting of 15 s ON and 30 s OFF. During the entire procedure, the microcentrifuge tubes containing the cells were kept in ice-cold water. Subsequently, ruptured cells were treated with 1 U μl^−1^ OmniCleave endonuclease (Lucigen) for 15 min at 37°C to release the DNA-bound protein pool. Cells were sonicated for two more cycles of 15 s ON and 30 s OFF. Finally, extract was clarified by centrifugation at 16,000 *g* for 15 min at 4°C, the supernatant fraction was collected, and the GFP concentration was estimated by measuring fluorescence in the microscopy chamber, as described in (Chakraborty et al. 2018). Extracts were used immediately in reconstitution assays, and the remainder was discarded after 5 hrs.

### Protein purification

Tubulin was purified from cow brains by thermal cycling and chromatography (Miller et al. 2010), and then labeled with HiLyte647 (Hyman et al. 1991). Human Bonsai and Broccoli Ndc80-GFP protein complexes were expressed in *Escherichia coli* and purified, as in (Zaytsev et al. 2015) and (Schmidt et al. 2012), respectively. Human Bronsai Ndc80-GFP construct (Wimbish et al. 2020) was generously provided for this work by J. DeLuca (Colorado State University). This construct contains the N-terminal fragment of Hec1 (1–506 amino acids (aa)) fused to a C-terminal fragment of the Spc25 (118–224 aa) and the Nuf2 protein (1–348 aa) fused to a C-terminal fragment of Spc24 (122–197 aa) (Wimbish et al. 2020). In this construct, the GFP-tag is located on the C-terminus of Hec1-Spc25 chain (Supplementary Figure 1). Full length wild type CENP-T-His or phospho-mimetic CENP-T-His and the untagged CENP-W genes were co-expressed in *Escherichia coli* and purified, as in (Gascoigne et al. 2011). Phospho-mimetic CENP-T mutant contains three residues mutated to aspartic acid (T11D, T27D, and T85D) in CDK phosphorylation sites.

### Preparation of antibody-coated beads

COOH-activated glass microbeads (0.5 μm, Bangs Labs) were coated with neutravidin (Thermo Fisher Scientific), as in (Grishchuk et al. 2008). Kinetochore proteins were conjugated to these beads using antibodies that recognize specific proteins or their tags: biotinylated anti-His-tag antibodies (6 µg ml^−1^, Qiagen), biotinylated anti-GFP antibodies (20 µg ml^−1^, Abcam), or biotinylated anti-rabbit antibodies (30 µg ml^−1^, Jackson ImmunoResearch). Antibodies at the indicated concentrations were incubated overnight with 4 mg ml^−1^ neutravidin beads, blocked with 2 mM biotinylated dPEG (2.5 kDa, Quanta BioDesign), washed extensively by centrifuging three times at 2,000 *g*, and resuspended in PBS-BSA buffer (135 mM NaCl, 2.5 mM KCl, 10 mM Na_2_HPO_4_, 1.8 mM KH_2_PO_4_ pH 7.2, 4 mg ml^−1^ BSA and 2 mM DTT). Subsequently, beads with anti-rabbit antibodies were coated with 50 µg ml^−1^ rabbit anti-CENP-T antibodies (Abcam) or 120 µg ml^−1^ rabbit anti-CENP-C antibodies (custom-made by Covance and affinity-purified in-house (Bassett et al. 2010)). A mixture of 35 µg ml^−1^ of anti-CENP-C antibodies and 25 µg ml^−1^ of anti-CENP-T antibodies were used to prepare beads for simultaneous recruitment of both CENP-C and CENP-T. Beads were incubated overnight with these antibodies at the indicated concentrations and washed three times. Antibody-coated beads were stored at 4°C for 7–10 days. Before usage, beads were sonicated for 10 s to reduce clumping.

### Preparation of CENP-T/W–coated beads

To immobilize recombinant CENP-T/W on the beads, 90 μl of 100-125 nM CENP-T/W (wild type or phospho-mimetic) in PBS-BSA buffer was mixed with 10 μl of beads coated with anti-CENP-T antibodies and incubated for 1.5–2 h at 4°C. Next, beads were washed three times, resuspended in 10 µl of the same buffer, and used immediately in bead-based reconstitution assays. We also immobilized recombinant CENP-T/W via its His-tag on the surface of the beads coated with anti-His-tag antibodies. These methods yielded similar levels of recruitment of Ndc80 and Mis12 complexes, so the resultant data were combined.

### Preparation of nucleosomes and their conjugation to beads

Human CENP-A and H3 nucleosomes were prepared using α-satellite sequence (147 bp) from human X-chromosome DNA (Figure 5A), as in (Sekulic et al. 2016). Briefly, Cy5-labelled 147-bp DNA was PCR-amplified with Cy5 custom primers in 96-well plates, followed by ResQ column purification. Sub-nucleosomal histone complexes were produced in *Escherichia coli*, purified, and assembled with DNA at equimolar ratio using gradual salt dialysis followed by thermal shifting for 2 h at 55°C (Figure 5B). Formation of nucleosomes was confirmed by 5% native PAGE analysis (Figure 5C). The nucleosomes were stored at 4°C for 1–2 months without loss of activity. To conjugate nucleosomes to beads, 10 µl of anti-His-tag antibody–coated beads were incubated for 1.5–2 h at 4°C with 100–200 nM of CENP-A nucleosomes, H3 nucleosomes, or a 1:1 mixture in PBS-BSA buffer supplemented with 4 mM MgCl_2_. Beads were washed three times, resuspended in 10 µl of the same buffer, and used immediately in bead-based reconstitution assays.

### Bead-based reconstitution assay

For the reconstitution assay with mitotic cell extract, 90 μl of freshly prepared extract was mixed with 10 μl of beads coated with antibodies, recombinant CENP-T/W, or nucleosomes, and incubated for 1 h at room temperature. To test the effect of the temperature, some samples were incubated at 4°C for 2–4 h. Lower temperature resulted in a similar level of kinetochore protein recruitment, so the resultant data were combined. After incubation with extract, beads were washed twice in lysis buffer supplemented with protease inhibitors, phosphatase inhibitors, and an ATP regeneration system. Beads were resuspended in 25 μl of PBS-BSA buffer supplemented 0.1 mg mL^−1^ glucose oxidase, 20 μg mL^−1^ catalase, 6 mg ml^−1^ glucose, and 0.5% β-mercaptoethanol. In experiments with nucleosomes, all incubations were carried out in buffers supplemented with 4 mM MgCl_2_ to avoid nucleosome unwrapping.

Purified recombinant proteins were used at the following concentrations, which were determined via the fluorescence microscopy approach (Chakraborty et al. 2018): 100–130 nM of Bonsai Ndc80-GFP, 120–130 nM Broccoli Ndc80-GFP, or 50–90 nM Bronsai Ndc80-GFP complexes. Incubation of these recombinant proteins with beads were carried out similarly in PBS-BSA; in Lysis Buffer supplemented with protease inhibitors, phosphatase inhibitors, and an ATP regeneration system; or in unlabeled mitotic extract prepared from control HeLa cells expressing no GFP-fused proteins.

After incubation with mitotic cell extract or recombinant proteins, beads were washed thoroughly, and 10 μl of beads were added on top of a coverslip, which was then covered with a glass slide to create a “wet” chamber. The chamber was sealed with VALAP (1:1:1 Vaseline/lanolin/paraffin), and images of the beads were taken immediately under the microscope in bright field and fluorescence channels. To quantify recruitment of the fluorescently labeled kinetochore complexes, images were analyzed using the custom-made MATLAB program “Quantification of bead brightness” (available on the Grishchuk Lab website (med.upenn.edu/grishchuklab/protocols-software.html)). This program measures the integral fluorescence intensity of beads selected using bright field images. Brightness of the same size area located near each bead is subtracted to minimize variability in background intensity. Typically, 50–100 beads were analyzed for each independent experiment, and average fluorescence brightness was calculated.

### Bead-based microtubule binding assay

Stabilized fluorescent microtubules were prepared as described in (Chakraborty et al. 2018) from a mixture of unlabeled and HiLyte647-labeled tubulin (5:1, total tubulin concentration 100 μM) and 1 mM GMPCPP (Jena Bioscience, Jena, Germany) incubated at 37°C for 20 min. A flow chamber was prepared with a silanized coverslip and a regular glass slide using spacers made from two strips of double-sided tape, as in (Chakraborty et al. 2018). The surface of the coverslip was activated by incubation with 22.5 μM of biotin-BSA (Sigma-Aldrich) for 10 min, followed by incubation for 10 min with 25 μM neutravidin. Next, beads coated with kinetochore complexes via biotinylated antibodies were diluted five times in PBS-BSA buffer, and 25 μl of beads were introduced to the flow chamber. The chamber was incubated for 10 min to allow immobilization of beads onto the neutravidin-coated coverslip. The chamber was blocked with 1% Pluronic F-127 (Sigma-Aldrich) and 0.1 mM biotin-PEG and incubated for 15 min with GMPCPP-stabilized HiLyte647-labeled microtubules diluted 1:100 in imaging buffer (BRB80: K-PIPES 80 mM, pH 6.9, 4 mM Mg_2_, 1 mM EGTA, supplemented with 4 mg ml^−1^ BSA, 2 mM DTT, 0.1 mg ml^−1^ glucose oxidase, 20 μg ml^−1^ catalase, 6 mg ml^−1^ glucose, 0.5% β-mercaptoethanol). Imaging was performed on a Nikon Eclipse Ti-E inverted microscope as described in “Immunofluorescence of cultured cells”. Z-stacks were taken in a ±0.25 μm range with 0.05-μm steps in the DIC and fluorescence channels to visualize beads and microtubules, respectively. At least four imaging fields per chamber were collected and analyzed using the Fiji software (Schindelin et al. 2012). The number of immobilized beads and the number of microtubules attached to these beads were counted for each imaging field. Efficiency of microtubule binding was calculated as the total number of microtubules attached to the beads normalized on the total number of beads in each field.

### TIRF microtubule binding assay

Binding of GFP-labeled protein complexes to microtubules *in vitro* was analyzed using TIRF microscopy. Custom-made flow chambers were assembled with silanized coverslips (22×22 mm), and solutions were exchanged with a peristaltic pump as in (Volkov et al. 2014). To immobilize microtubules, the coverslip of the flow chamber was coated with anti-tubulin antibodies (Serotec) and blocked with 1% Pluronic F-127, and then fluorescently labeled microtubules were flowed into the chamber. Freshly prepared cell extract was diluted in imaging buffer to achieve a 2.5–3 nM concentration of the GFP-labeled component. The mixture was perfused continuously into the chamber using a peristaltic pump at 15 µl min^-1^ at 32°C. Experiments with purified Bronsai Ndc80-GFP complexes were carried out similarly using an aliquot of purified protein, which was thawed and centrifuged at 157,000 *g* for 15 min at 4°C to remove aggregates. To account for possible stage drift, images of microtubules were taken at the beginning and end of each 30-s imaging sequence in GFP-channel. GFP images were collected in TIRF mode using stream acquisition with 100-ms exposures and the 488 nm laser at 10% power. An Andor iXon3 EMCCD camera was used with 5x conversion gain, 999 EM Gain Multiplier, 10 MHz readout speed, and 14-bit sensor mode. Kymographs were assembled using the Fiji software. To quantify microtubule decoration with various GFP-fused kinetochore complexes, microtubule area was selected on the kymograph plot in HiLyte647-channel. Average GFP fluorescence intensity was calculated in this area and the area located near each microtubule. To calculate normalized GFP brightness, the background value was subtracted, and the difference was normalized against the concentration of soluble GFP-labeled protein.

## Acknowledgements

We are grateful to Dr. M. Lampson and Dr. H. Zhang (University of Pennsylvania) for providing HeLa cell lines stably expressing the GFP-fused kinetochore proteins CENP-C, Mis12, Nuf2, Spc24, Spc25, and CENP-T. We thank Dr. D.W. Cleveland (University of San Diego) for providing DLD-1 Flp-In T-REx cells stably expressing CENP-C^AID-EYFP/AID-EYFP^. Purified Bronsai Ndc80 protein was generously provided by Dr. J. DeLuca (Colorado State University). CENP-T/W protein was purified by Florencia Rago. We appreciate the technical assistance and insightful suggestions of Ricardo Sousa, Eirini Zoupou, and Aaron (Po-Tao) Chen. We thank Dr. J. Dawicki-McKenna and other members of Grishchuk, Black, and Lampson labs for sharing reagents and for helpful discussions.

## Author contributions

E.V.T. performed all experiments on kinetochore assembly *in vitro*. P.K.A. reconstituted and characterized fluorescently labeled recombinant nucleosomes and generated the DLD-1 cell line expressing GFP-Dsn1-SD. I.M.C. provided recombinant CENP-T/W and the HeLa cell line expressing GFP-Dsn1-Δ91-113, and contributed to the design of the research. E.V.T., B.E.B., P.K.A., and E.L.G. devised experiments investigating the role of the nucleosome-dependent assembly pathway. E.V.T. and E.L.G. designed the research project, analyzed the data, and wrote the paper with input from B.E.B., I.M.C., and P.K.A.

## Abbreviations list

BSA: bovine serum albumin
CCAN: constitutive centromere-associated network
CENP: centromere protein
DAPI: 4′,6-diamidino-2-phenylindole
DIC: differential interference contrast
PBS: phosphate-buffered saline
SD: standard deviation
SEM: standard error of the mean
TIRF: total internal reflection fluorescence.

