## Supplementary Material for "Reconstitution of human kinetochore in mitotic cell extracts reveals permitted and restricted assembly steps"

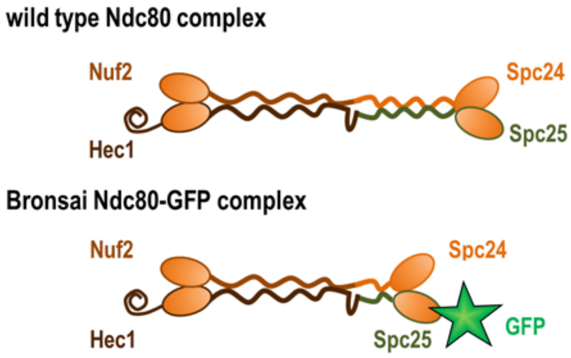


**Supplementary Figure 1. Recombinant Ndc80 constructs.** Illustration of the wild-type and mutant GFP-labeled Ndc80 constructs (not to scale) used in this study.

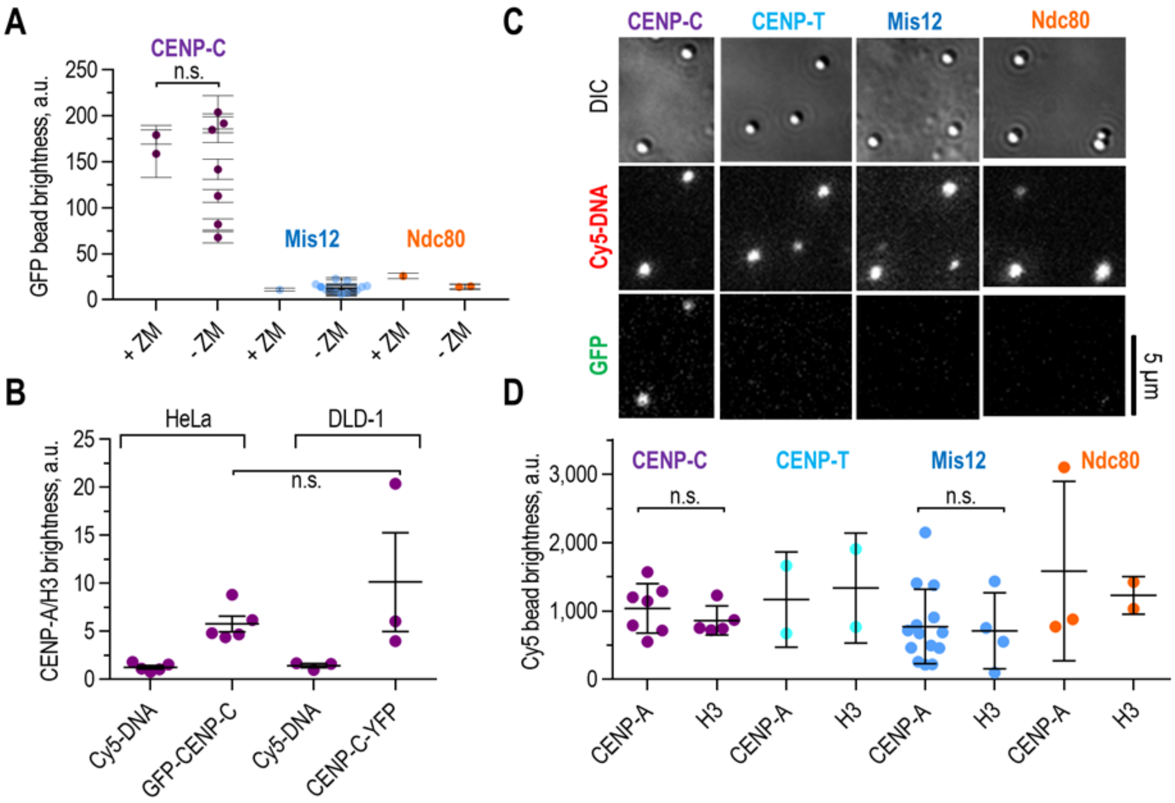


**Supplementary Figure 2. Reconstitution of kinetochore complexes on CENP-A nucleosomes.** (A) Average GFP brightness of CENP-A–coated beads after incubation with mitotic cell extracts containing either GFP-CENP-C, Mis12-GFP, or Ndc80-GFP complexes. Extracts were prepared from HeLa cells grown in the presence or absence of 2.5 µM ZM447439, an Aurora B inhibitor, (for 1 h before harvesting. Here, data show means with SEM; each point represents an independent experiment. (B) The ratio of CENP-A– and H3-coated bead brightness in the Cy5-DNA and GFP (YFP)-channels, showing that in both HeLa and DLD-1 cell extracts CENP-A, but not H3, recruits CENP-C. Error bars indicate SEM. On panels (A) and (B), data for HeLa cells are the same as shown on Figure 5F,G. (C) Representative images of the beads in DIC and fluorescence channels, showing CENP-A nucleosomes with Cy5-DNA and level of recruited GFP-fused kinetochore proteins (CENP-C, CENP-T, Mis12, and Ndc80 complexes). (D) Average DNA brightness of beads shown on Figure 5G, which indicate similar densities of nucleosome coating. Error bars indicate SD. P-values were calculated by unpaired t-test: n.s., $p > 0.05$. For more detailed statistics, see Source data.

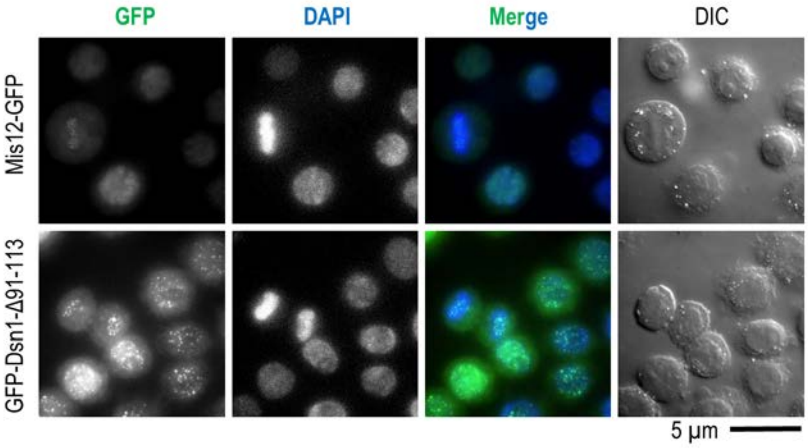


**Supplementary Figure 3. Characterization of GFP-Dsn1-Δ91-113 HeLa cell line.** Representative images of asynchronous HeLa cells expressing either Mis12-GFP or GFP-Dsn1-Δ91-113 mutant. DNA was stained with DAPI. Images illustrate constitutive recruitment of GFP-Dsn1-Δ91-113 to kinetochores. Kinetochore localization of GFP-Dsn1-Δ91-113 was observed in 100% of interphase cells (N=148 cells), whereas wild type Mis12-GFP was found in kinetochores of 52% of interphase cells (N=114 cells).

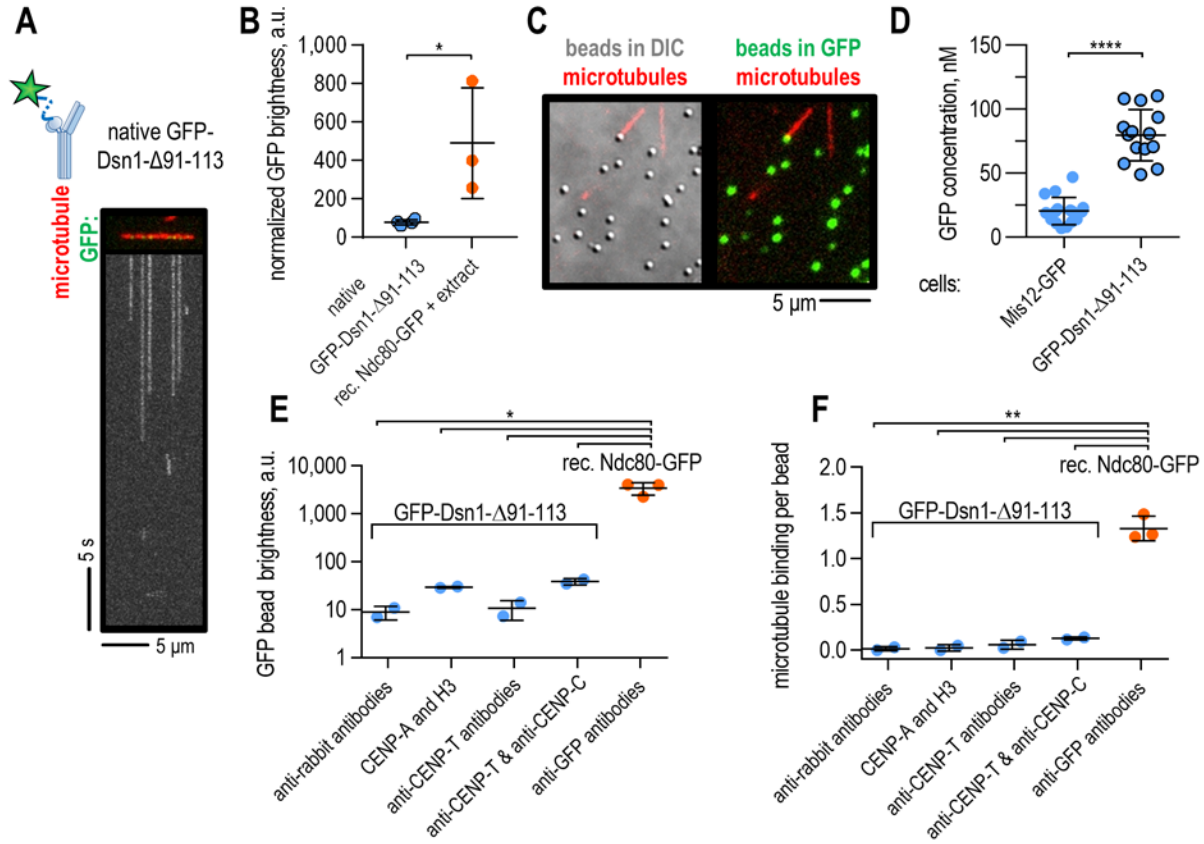


**Supplementary Figure 4. Reconstitution of kinetochore assembly on different nucleators using autoinhibition-deficient Mis12 complex.** (A) Representative image showing a microtubule (red) and GFP-fused Dsn1-Δ91-113 complexes (green); mobility of complexes over 30 s is shown on kymographs below. (B) Average GFP brightness of microtubule decoration over 30 s, quantified based on kymographs and normalized against GFP concentration; data for recombinant Ndc80 complex with unlabeled extract is the same as in Figure 2C, and is provided for illustrative purposes. Here and in other graphs in this figure, means are shown with SD, and each point represents an independent experiment. P-values were calculated by unpaired t-test: *, $p<0.05$; **, $p<0.01$; ****, $p<0.0001$. For more detailed statistics, see Source data. (C) Representative image showing GFP-fused Dsn1-Δ91-113 complexes clustered on beads, which were immobilized on the coverslip and exposed to floating microtubules (red). Beads are shown in the DIC (left) and in GFP channels (right). (D) Concentrations of GFP-fused Mis12 or Dsn1-Δ91-113 protein complexes in mitotic HeLa cell extracts, estimated by fluorescence measurement; data regarding the Mis12 complex are the same as in Figure 1E. (E) GFP bead brightness showing recruitment of GFP-fused Dsn1-Δ91-113 to beads via different tethers. (F) Average number of bead-bound microtubules, normalized against the number of beads per imaging field. In panels (E and F), as a positive control, data for beads coated with 90 nM recombinant Ndc80 complex in the presence of unlabeled mitotic extract are shown in orange; these data are the same as in Figure 3C,D.

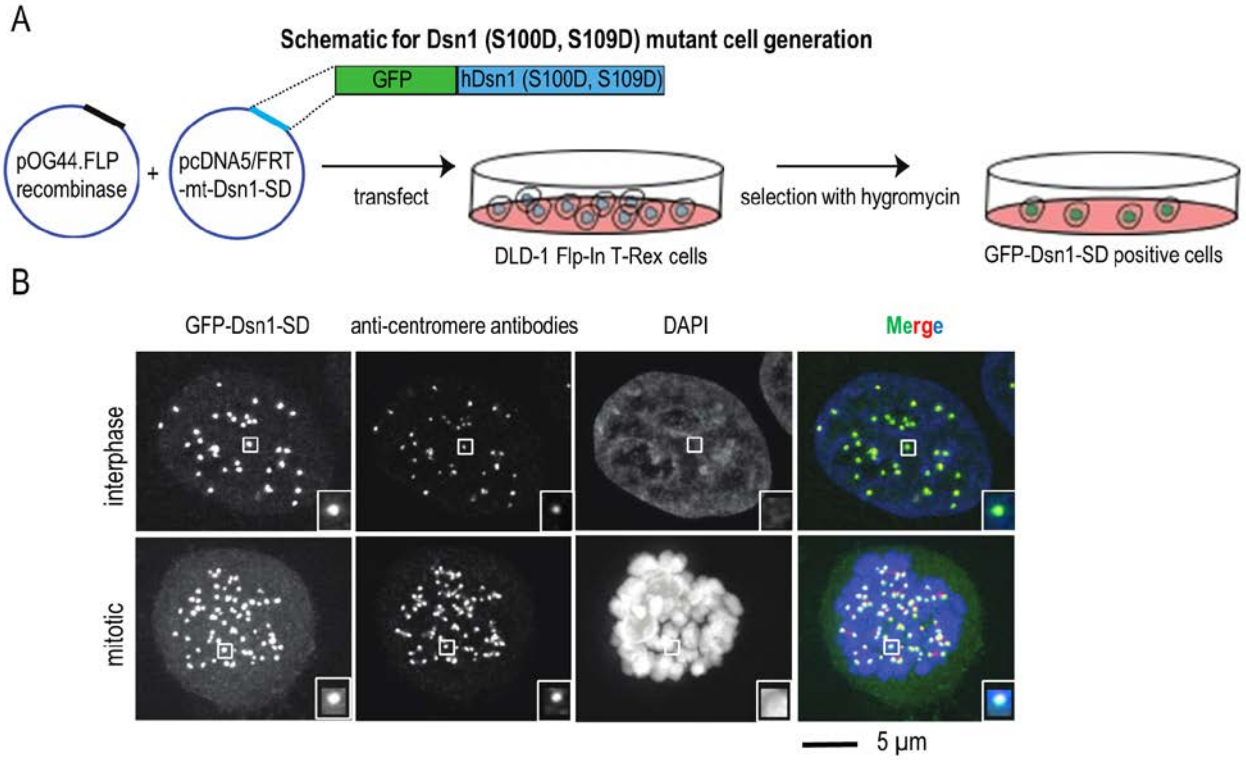


**Supplementary Figure 5. Generation of the phospho-mimetic mutant GFP-Dsn1-SD in the DLD-1 cell line.** (A) Schematic of generation of the DLD-1 Flp-In T-REx cell line expressing GFP-Dsn1 (S100D, S109D) at the FRT site. Plasmids containing Dsn1-SD and FLP recombinase were transfected into DLD-Flp-In T-REx cells, and hygromycin selection was performed for 10 days, followed by analysis of Dsn1-SD localization in hygromycin-resistant colonies. (B) Representative images of interphase (asynchronous) and mitotic (S-Trityl-L-cysteine treated) cells expressing GFP-Dsn1-SD mutant. Kinetochores were stained with anti-centromere antibodies and DNA with DAPI.

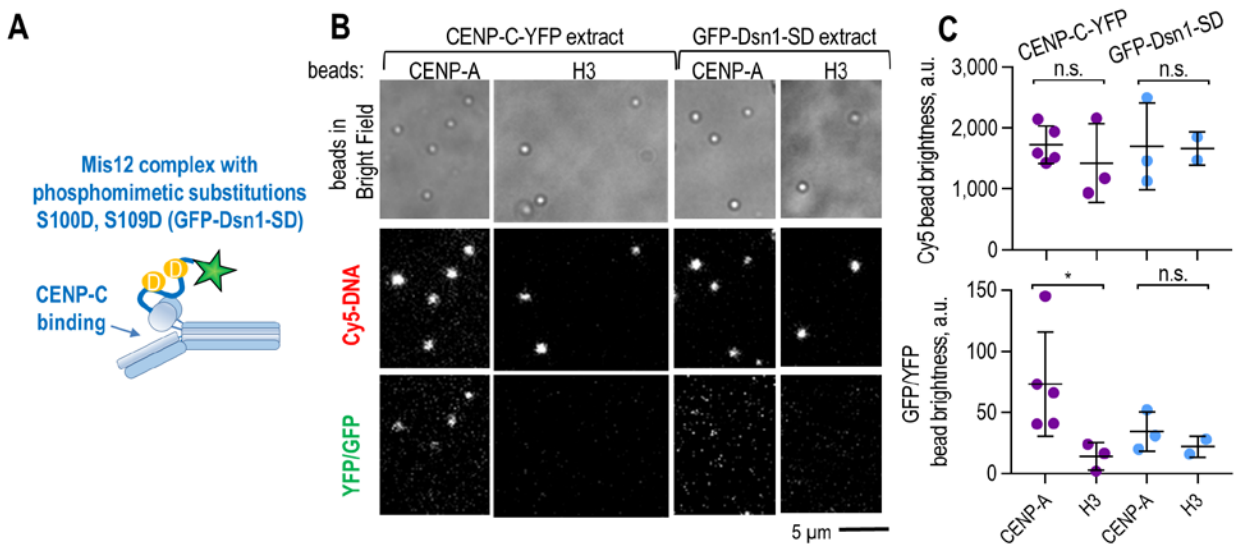


**Supplementary Figure 6. Lack of significant recruitment of the phospho-mimetic Mis12 complex from DLD-1 cell extract.** (A) Scheme showing mutant Mis12 complex carrying phospho-mimetic substitutions S100D and S109D in the Dsn1 subunit. (B) Representative images of beads in DIC and fluorescence channels, showing nucleosomes with Cy5-DNA and CENP-C-YFP or GFP-Dsn1-SD. YFP and GFP signals were captured at the same settings. (C) Average brightness of beads showing levels of associated DNA (top) and GFP (bottom), corresponding to the densities of nucleosome and either CENP-C-YFP or GFP-Dsn1-SD, respectively (means are shown with SD; each point is an independent experiment). Although YFP and GFP brightness cannot be compared directly, note the significant difference in the level of CENP-C-YFP recruitment to CENP-A vs. H3 nucleosomes. Difference in GFP-Dsn1-SD recruitment to CENP-A vs. H3 nucleosomes is not significant. P-values were calculated by unpaired t-test: n.s., $p>0.05$; *, $p<0.05$. For more detailed statistics, see Source data.
